# Evolutionary modes of *wtf* meiotic driver genes in *Schizosaccharomyces pombe*

**DOI:** 10.1101/2024.05.30.596636

**Authors:** Yan-Hui Xu, Fang Suo, Xiao-Ran Zhang, Tong-Yang Du, Yu Hua, Guo-Song Jia, Jin-Xin Zheng, Li-Lin Du

## Abstract

Killer meiotic drivers (KMDs) are a class of selfish genetic elements that defy Mendel’s law and bias transmission in their favor by destroying meiotic progeny that do not carry them. How KMDs evolve is not well understood. In the fission yeast *Schizosaccharomyces pombe*, the largest gene family, known as the *wtf* genes, is a KMD family that causes intraspecific hybrid sterility. Here, we investigate how *wtf* genes evolve using long-read-based genome assemblies of 31 distinct *S. pombe* natural isolates, which encompass the known genetic diversity of *S. pombe*. Our analysis, involving nearly 1,000 *wtf* genes in these isolates, yields a comprehensive portrayal of the intraspecific diversity of *wtf* genes. Leveraging single-nucleotide polymorphisms in adjacent unique sequences, we pinpoint *wtf*-gene-containing loci that have recently undergone gene conversion events and infer their pre-gene-conversion state. These events include the revival of *wtf* pseudogenes, lending support to the notion that gene conversion plays a role in preserving this gene family from extinction. Moreover, our investigation reveals that solo long terminal repeats (LTRs) of retrotransposons, frequently found near *wtf* genes, can act as recombination arms, influencing the upstream regulatory sequences of *wtf* genes. Additionally, our exploration of the outer boundaries of *wtf* genes uncovers a previously unrecognized type of directly oriented repeats flanking *wtf* genes. These repeats may have facilitated the early expansion of the *wtf* gene family in *S. pombe*. Our findings enhance the understanding of the mechanisms influencing the evolution of this KMD gene family.

## Introduction

Under the framework of Darwinian evolution, natural selection governs the evolution of genes. When one allele of a gene confers greater benefits to the organism compared to another allele, it secures a selective advantage. The fair competition among gene alleles is ensured by Mendel’s law, which dictates that during meiosis, two alleles of a gene have an equal chance to be inherited by the next generation. However, not all genes obey Mendel’s law. Killer meiotic drivers (KMDs) are a category of selfish genetic elements that promote their own propagation at the expense of the organism by destroying meiotic products that do not carry them (Bravo Núñez *et al*. 2018b). KMDs have been identified in a wide range of eukaryotic species, including plants, animals, and fungi (Burga *et al*. 2020; Zanders and Johannesson 2021; Vogan *et al*. 2022; Winkler and Lindholm 2022; Saupe and Johannesson 2022; Wang *et al*. 2023, 2024; Lai and Vogan 2023). A common strategy employed by KMDs to selectively kill meiotic products is through expressing both a “toxin” product and an “antidote” product. While the toxin can affect all meiotic products, only those carrying the KMD are protected by the antidote.

In the unicellular model eukaryotic organism the fission yeast *Schizosaccharomyces pombe*, *wtf* genes—named for their frequent association with the solo long terminal repeats (LTRs) of Tf retrotransposons (*wtf* stands for with Tf LTRs)—have been identified as a KMD gene family that causes low spore viability during outcrossing (Zanders *et al*. 2014; Hu *et al*. 2017; Nuckolls *et al*. 2017). An active *wtf* driver gene can produce two overlapping protein isoforms through alternative transcription initiation (Nuckolls *et al*. 2017). The long isoform, which contains the amino acid sequence encoded by exon 1, functions as the antidote, while the short isoform, with its transcript initiated from within intron 1, acts as the toxin. Both the antidote and the toxin are multi-transmembrane proteins, with the antidote containing a longer N-terminal cytosolic tail. The cytosolic tail of the antidote contains multiple PY motifs (Leu/Pro-Pro-X-Tyr), which mediate the ubiquitination-dependent sorting of the antidote from the Golgi to the endosome (Zheng *et al*. 2023). This sorting mechanism serves two purposes. Firstly, it prevents the antidote from exerting toxicity. Secondly, it enables the antidote, which can interact with the toxin, to neutralize the toxin by altering its trafficking route.

In the reference *S. pombe* genome—that of an isolate from French grape juice, there are 25 *wtf* genes designated as *wtf1* to *wtf25* based on their genomic positions (Wood *et al*. 2002; Bowen *et al*. 2003). These 25 *wtf* genes reside in 20 genomic loci, including 15 loci each containing a single *wtf* gene and 5 loci each containing a pair of tandemly oriented *wtf* genes. Four of the 25 genes—*wtf7*, *wtf11*, *wtf14*, and *wtf15*— are highly divergent from each other and the remaining *wtf* genes (Hu *et al*. 2017; Eickbush *et al*. 2019). We hereafter refer to them as non-typical *wtf* genes. These non-typical *wtf* genes do not exhibit meiotic driver activities (Bravo Núñez *et al*. 2020). The remaining 21 wtf genes, which are considered typical *wtf* genes, show a higher level of relatedness. Within this group, there are 4 genes that encode both toxin and antidote, and therefore have the potential to act as meiotic drivers. These genes will be referred to as 2-isoform genes. Additionally, there are 8 genes that encode only antidote, referred to as antidote-only genes, and 9 pseudogenes (Hu *et al*. 2017; Eickbush *et al*. 2019).

It has been theorized that a toxin-antidote meiotic driver undergoes a two-phase decay once it becomes fixed in a population (López Hernández and Zanders 2018). Initially, it transitions into an antidote-only variant, which does not suffer a competitive disadvantage against the active driver allele. Eventually, after the fixation of the antidote-only variant, it then transitions into a pseudogene. However, in the case of *wtf* genes, the antidote-only genes are not merely decaying intermediates; they can actually function as suppressors of closely related active 2-isoform genes located elsewhere in the genome (Bravo Núñez *et al*. 2018a). Furthermore, antidote-only *wtf* genes pose an ongoing threat to the host due to the inherent toxicity of the antidote, necessitating the host’s ubiquitination machinery to prevent the release of this hidden toxicity (Zheng *et al*. 2023).

Two previous studies have obtained the sequences of the complete set of *wtf* genes in 3 additional *S. pombe* isolates: JB1180 (also known as *S. kambucha*, abbreviated *Sk*), JB916/FY29033, and JB4/CBS5557 (Hu *et al*. 2017; Eickbush *et al*. 2019). By comparing the *wtf* genes in different isolates, we and the Zanders group found that *wtf* genes undergo rapid evolution through several types of changes, including alterations in gene sequence via non-allelic gene conversion (Hu *et al*. 2017; Eickbush *et al*. 2019), expansion and retraction of two types of intragenic repeats—a 33-bp repeat and a 21-bp repeat (Eickbush *et al*. 2019), and variations in gene number within a specific locus (Hu *et al*. 2017; Eickbush *et al*. 2019). However, the exact processes underlying these changes remain unknown, as no previous attempts have been made to infer the pre-alteration state. Furthermore, these 4 isolates may not adequately represent the *S. pombe* species, given the existence of approximately 60 distinct natural isolates of *S. pombe* (Jeffares *et al*. 2015; Tao *et al*. 2019; Tusso *et al*. 2022). Genomic analysis conducted on this broader set of isolates has identified two ancient lineages that are estimated to have diverged around 80,000 years ago, which have only recently experienced admixture (Tao *et al*. 2019; Tusso *et al*. 2019). The implications of this admixture on the evolution of *wtf* genes remain unknown.

In collaboration with the Zanders group, we have recently shown that *wtf* genes are not only present in *S. pombe*, but also in three other fission yeast species (*S. octosporus*, *S. osmophilus*, and *S. cryophilus*) that diverged from *S. pombe* approximately 100 million years ago (De Carvalho *et al*. 2022). Since some of the *wtf* genes in *S. octosporus* are active drivers, it is likely that the ancestors of *wtf* genes existing 100 million years ago are KMD genes. This is surprising, as KMDs are thought to unavoidably undergo decay after fixation and therefore have short evolutionary lifespans. How this KMD family has managed to persist for such a long period of time without becoming extinct is a fascinating question. One possibility is that the decay of wtf genes is not irreversible and a *wtf* pseudogene can potentially be converted back into an active gene. However, there is currently no direct evidence indicating that the revival of *wtf* pseudogenes can actually occur.

In *S. octosporus* and *S. osmophilus*, most of the *wtf* genes are flanked by direct repeats of 5S rDNA genes (De Carvalho *et al*. 2022). In these two species, there are species-specific 5S rDNA-flanked *wtf* genes that show synteny with *wtf*-free regions that contain a single 5S rDNA gene in the other species. This suggests that *wtf* genes have expanded into new genomic locations through duplication into 5S rDNA genes that are dispersed throughout the genome. In *S. pombe*, *wtf* genes are not associated with 5S rDNA genes. Instead, they are associated with a different type of dispersed repeats—the solo LTRs of retrotransposons. Whether LTRs mediate the spread of *wtf* genes to new locations in *S. pombe* remains an open question. Furthermore, the potential impact of LTRs on *wtf* genes in other ways has not yet been investigated.

In this study, we comprehensively identify and analyze the *wtf* genes in long-read-based genome assemblies of 31 *S. pombe* natural isolates that represent the full genetic diversity of this species. This extensive dataset enables us to infer the nature of recent evolutionary events affecting *wtf* genes. Among the inferred gene conversion events, we identify several that result in the revival of *wtf* pseudogenes. Furthermore, we have discovered a role of *wtf*-adjacent solo LTRs in facilitating gene conversion. These conversions have led to alterations in the upstream regulatory sequences of *wtf* genes. Moreover, we identify previously unknown direct repeats that flank *wtf* genes, which may have contributed to the early spread of *wtf* genes in *S. pombe*.

## Materials and Methods

### Long-read PacBio sequencing of 20 *S. pombe* isolates

A previously published study of 161 *S. pombe* isolates (referred to as JB strains hereafter because their strain names all begin with the initials JB) identified 57 clades, each differing from the others by at least 1,900 SNPs (Jeffares *et al*. 2015). Within each clade, if there were multiple strains, any two strains were found to differ by less than 150 SNPs, indicating clonality or near-clonality. The study selected a set of 57 JB strains, known as “non-clonal strains”, to represent these 57 clades. In our study, we obtained PacBio long-read sequencing data for 20 *S. pombe* strains, with 18 of them corresponding to 18 clades previously identified by Jeffares et al.

Out of the 18 strains, 12 were JB strains acquired from the Yeast Genetic Resource Center (YGRC) of the National BioResource Project (NBRP) in Japan. These JB strains have been deposited at YGRC/NBRP by the Bähler laboratory. Among the other strains, one was a CRISPR-engineered heterothallic derivative of JB938 (Zhang *et al*. 2018). Two strains were heterothallic derivatives of CBS5557 and CBS5682, which correspond to JB4 and JB874, respectively. Another strain, CBS356, belongs to the clade represented by the non-clonal strain JB864. One strain, DY29153, was obtained from the China General Microbiological Culture Collection Center (CGMCC) and belongs to the clade represented by the non-clonal strain JB929. We also included a laboratory strain, DY38751 (*h+ mat1PΔ17 leu1-32 lys1-131 ade6-M216 ura4-D18*), which belongs to the clade represented by the non-clonal strain JB22. For simplicity, the above-mentioned 18 strains will be referred to by the names of the non-clonal strains that represent the corresponding clades.

In addition to these 18 strains, two others analyzed in this study, DY34373 and DY39827, were obtained from CGMCC and the US Department of Agriculture Agricultural Research Service Culture Collection (NRRL), respectively, and their genomes differ from those of the JB strains (Tao *et al*. 2019; Tusso *et al*. 2022). The PacBio long-read sequencing of the 20 strains was performed using the RS II or the Sequel platform. Illumina short-read sequencing data were also obtained. The genome sequencing data used in this study have been deposited at the NCBI SRA database, with the accession numbers listed in Supplementary Table 1.

### De novo assembly of the nuclear genomes of the 20 *S. pombe* isolates

To obtain high-quality genome assemblies and minimize the impact of assembler-specific errors, we employed three long-read assemblers (Canu, Flye, and Raven) for de novo genome assembly (Supplementary Fig. 1A). Genome assemblies were generated using Canu 1.8 (-pacbio-raw, genomeSize=12.5m, useGrid=false), Flye 2.6 (--pacbio-raw, --genome-size 12m, --thread 4), and Raven 0.07 (default parameter) (Koren *et al*. 2017; Kolmogorov *et al*. 2019; Vaser and Šikić 2021). The assemblies underwent one round of polishing using long-read data and the polishing tool GCpp 0.0.1, followed by three rounds of polishing using short-read data and the polishing tool Pilon 1.23 (Supplementary Fig. 1A).

Next, we used short-read data to filter out contaminating contigs and contigs corresponding to the mitochondrial genome. First, we removed short reads that could be mapped to the corresponding mitochondrial genomes (Tao *et al*. 2019). Then, we mapped the remaining short reads (nuclear genome reads) to the corresponding assembly. If a contig had more than half of its length covered by nuclear genome reads with depths lower than one-tenth of the average depth of the entire assembly, it was removed from the assembly. We also eliminated all contigs shorter than 1 kb. The nuclear genome assemblies have been deposited at the NCBI Assembly database (accession numbers are listed in Supplementary Table 1).

### Assembly quality assessment

The completeness, continuity, and correctness of our 60 genome assemblies (3 assemblies per isolate) were assessed using QUAST, BUSCO, and asmgene, and by mapping short-read data to the assemblies (Supplementary Fig. 1A) (Gurevich *et al*. 2013; Simão *et al*. 2015; Cheng *et al*. 2021). Analysis using QUAST 5.0.2 showed that the median N50 values are 1.69 Mb, 1.11 Mb, and 1.17 Mb for the 20 assemblies generated by Raven, the 20 assemblies generated by Flye, and the 20 assemblies generated by Canu (Supplementary Fig. 1B). Analysis using BUSCO 3.1.0 showed that, out of 1315 ascomycota_odb9 BUSCO orthologs, all assemblies except JB1180-Canu and JB879-Canu have fewer than 60 missing BUSCO orthologs (as a control, the *S. pombe* reference genome has 41 missing BUSCO orthologs) (Supplementary Fig. 1C). Using Minimap2’s paftools.js (version: 2.17-r941) asmgene script (with a sequence similarity cutoff at 95%), we found that, except for JB1180-Canu and JB879-Canu, all assemblies contain completely assembled CDSs for more than 97% of the 5152 protein-coding genes in the *S. pombe* reference genome (Supplementary Fig. 1D).

We mapped short-read data to the assemblies using BWA and called variants using Samtools (version: 0.1.18), GATK (version: 4.1.4.1), and DeepVariant (version: 0.10.0) (Li and Durbin 2009; Li *et al*. 2009; McKenna *et al*. 2010; Poplin *et al*. 2018). The number of SNPs and indels supported by at least two variant callers was used as an indication of the correctness of the assemblies. All assemblies, except for JB873-Flye and DY34373-Flye, have fewer than 50 variants (Supplementary Fig. 1E).

### Previously published nuclear genome assemblies of 17 *S. pombe* isolates

PacBio sequencing-based nuclear genome assemblies of 17 *S. pombe* strains generated in a previous study were downloaded from the NCBI Assembly database (https://www.ncbi.nlm.nih.gov/assembly/organism/4896/all/) (Tusso *et al*. 2019). These assemblies were aligned to the *S. pombe* reference genome using minimap2 and sequence variants were called using paftools (Li 2018). SNP variants found in the assemblies were compared to SNP variants called from Illumina sequencing data (Jeffares *et al*. 2015). For 15 assemblies, SNP variants found in the assemblies matched those called from Illumina sequencing data. However, SNP variants found in the other two assemblies, EBC131_JB1171_v01_pb and EBC132_JB1174_v01_pb, did not match those called from the Illumina sequencing data of JB1171 and JB1174, respectively. Instead, SNP variants found in these two assemblies matched SNP variants called from the Illumina sequencing data of JB900. JB900 is the non-clonal strain representing a clade containing multiple JB strains. We hereafter refer to these two assemblies as JB900_EBC131 and JB900_EBC132.

### Identifying *wtf* genes in the genome assemblies

Studying the evolution of *wtf* genes requires the sequences of not only the *wtf* genes themselves but also the sequences of the genomic regions surrounding the *wtf* genes. To obtain *wtf* gene sequences together with flanking genomic sequences, we first searched the genome assemblies for the 24 genomic loci known to harbor *wtf* genes (Hu *et al*. 2017; Eickbush *et al*. 2019). Using the sequences of unique protein-coding genes flanking the *wtf* genes at these loci as queries, we performed BLASTN searches against the genome assemblies. If the unique gene on one side of a locus in the reference genome was not found within 25 kb of the unique gene on the other side of the locus in the reference genome, we manually examined the assembly to determine whether and how the synteny may have been disrupted by a genome rearrangement event. We identified 5 instances where genome rearrangement events had occurred within or near *wtf* genes, resulting in a change in their flanking genes (Supplementary Fig. 2).

Next, we identified the sequences of *wtf* genes at these loci by conducting BLASTN searches using the sequences of known *wtf* genes (from the start of the conserved_up region, an approximately 288-bp sequence upstream of the start codon of the antidote (Hu *et al*. 2017), to the stop codon of the coding sequence) as queries. After identifying *wtf* genes in the 24 genomic loci within a genome assembly, we masked the sequences of all identified *wtf* genes in the assembly and performed BLASTN searches against the *wtf*-masked assembly using the sequences of known *wtf* genes as queries. No additional *wtf* genes were found outside of the 24 genomic loci in any of the genome assemblies. For a consistent nomenclature of the *wtf* genes in different isolates, we assigned names to these 24 loci with the prefix *wts* (for *wtf* sites), followed by numbers. Please refer to the Results section for further details on the nomenclature. We note here that when there is more than one *wtf* gene at a locus, they are nearly always tandemly oriented, with the only exceptions being the *wts1* loci in JB758 and JB943.

### Determining the sequences of *wtf* genes in the genome assemblies

In the 20 isolates whose genomes were assembled in this study, we identified a total of 614 *wtf* genes. For 603 of these 614 *wtf* genes, no sequence differences exist between the assemblies generated by the three assemblers (Raven, Flye, and Canu). For the remaining 11 *wtf* genes, 10 have sequences that agree between two assemblies but differ from the third assembly. In these cases, we selected the *wtf* gene sequence supported by two assemblies. However, there is one gene (JB864-*wts28*) for which the exact sequence could not be determined because the three assemblies disagree with each other.

For 4 of these 20 isolates, JB4, JB760, JB873, and JB1206, genome assemblies were independently generated in another study (Tusso *et al*. 2019). For all but one gene (JB873_*wts0403*), the sequences of *wtf* genes in these 4 isolates determined based on the genome assemblies generated in this study agree with the sequences we extracted from the genome assemblies generated by Tusso et al. We used the JB873_*wts0403* sequence determined in this study for subsequent analyses. Genome assemblies of two different laboratory *S. pombe* strains, DY38751 and EBC2, were generated in this study and in Tusso et al., respectively. They are reference genome background strains. For all but one of the *wtf* genes in the reference genome, no differences were found between the sequences in the reference genome, the sequences determined from the assemblies of DY38751, and the sequences extracted from the assembly of EBC2. The exception is *wts0403a*, which showed the same sequence in the reference genome and in the EBC2 assembly, but showed a loss of one repeat unit in the 21-bp repeat region in the DY38751 assemblies. This is likely due to recent divergence in laboratory settings. We used the sequence of *wts0403a* in the reference genome for subsequent analyses. For the isolate JB1180 (also known as *S. kambucha*), the *wtf* gene sequences determined in this study agree with those previously reported in an independent study (Eickbush *et al*. 2019). The *wtf* gene sequences extracted from the JB900_EBC131 assembly are identical to those extracted from the JB900_EBC132 assembly.

In total, we obtained the sequences of 965 *wtf* genes and their flanking regions from 31 distinct *S. pombe* isolates. This includes the sequences in the reference genome background (with JB22 as the non-clonal strain) and the sequences extracted from the PacBio-based genome assemblies of 30 distinct *S. pombe* natural isolates (11 isolates with assemblies generated by Tusso et al., 15 isolates with assemblies generated in this study, and 4 isolates with assemblies generated by both Tusso et al. and this study). The *wtf* gene sequences, including the flanking regions, from these 31 isolates have been deposited in the NCBI GenBank (accession numbers listed in Supplementary Table 2). For our analysis of *wtf* gene evolution, we also included the sequences of *wtf* genes of the JB916 isolate (NBRP strain name FY29033) (Eickbush et al. 2019).

### PacBio-based amplicon sequencing

Eight genomic loci harboring *wtf* genes (*wts0403*, *wts8*, *wts9*, *wts13*, *wts1817*, *wts19*, *wts23*, *wts27*) were selected for PacBio-based amplicon sequencing. PCR primers specific for each locus were designed using the Primer3 software. The first-round PCR used locus-specific primers tailed with M13 forward and reverse primers, respectively. The resulting PCR products were used as templates for the second-round PCR, which used barcoded M13 primers to generate barcoded PCR products. PCR amplification was performed using PrimeSTAR GXL DNA polymerase (TaKaRa). The resulting barcoded PCR products were pooled for SMRTbell library construction and sequenced on the Sequel platform.

Amplicon sequencing reads were first demultiplexed using lima (version: 1.7.0). Consensus sequences from the amplicon sequencing reads were obtained using laa (version 2.4.2). In total, we obtained the sequences of 131 amplicons from *S. pombe* isolates whose clade affiliation differs from the 32 isolates mentioned above. The PacBio sequencing data for the amplicons have been deposited at NCBI under the BioProject accession PRJNA706838, and the amplicon sequences have been deposited at the NCBI GenBank (accession numbers listed in Supplementary Table 2).

### Gene structure annotation of *wtf* genes

The gene structures of *wtf* genes in the reference genome were annotated based on a published Iso-Seq dataset (Kuang *et al*. 2017). The annotated exons of *wtf* genes in the reference genome were then used as queries to perform BLASTN searches against the sequences of *wtf* genes from other genomes. The BLASTN results were used to annotate exons. To verify the boundaries of exons, sequence alignments were manually inspected.

### Classification of *wtf* genes

In the reference genome, 9 out of the 25 *wtf* genes are pseudogenes (Hu *et al*. 2017). We used a custom script to classify the *wtf* genes in the other genomes as intact genes or pseudogenes. For typical *wtf* genes (genes other than *wts7*, *wts1112a*, *wts14*, and *wts15*) that are intact, we classified them into two functional types: “2-isoform” and “antidote-only”. This classification was based on whether a gene has the two features associated with the toxin isoform: a 150-bp signature sequence in intron 1 (Hu *et al*. 2017) and an in-frame ATG start codon near the junction between intron 1 and exon 2. We identified the 150-bp signature sequence using BLASTN and the ATG start codon using a custom script. We observed that these two features were always either both present or both absent.

### Determining lineage ancestry of *wts* loci using SNPs in flanking sequences

All currently known *S. pombe* natural isolates are descended from two ancestral lineages referred to as the REF lineage (also known as the *Sp* lineage) and the NONREF lineage (also known as the *Sk* lineage) (Tao *et al*. 2019; Tusso *et al*. 2019). To determine the local lineage ancestry of *wts* loci in the 32 isolates, we examined the SNPs in the unique sequences surrounding each *wts* locus. Based on the SNP patterns, we classified a *wts* locus in an isolate as having either REF-lineage ancestry or NONREF-lineage ancestry. In 4 instances (JB1180_*wts5*, JB1110_*wts5*, JB918_*wts16*, and JB939_*wts1817*), where the upstream and downstream sequences of a *wts* locus belonged to different lineages, we considered the lineage ancestry of the locus to be uncertain.

### Analyzing the sequence diversity of *wtf* genes

To quantify the sequence diversity among syntenic *wtf* genes or among *wtf* genes within the same genome, we used the Sequence Demarcation Tool (SDT) to calculate pairwise identity of *wtf* gene sequences (Needleman-Wunsch algorithm as implemented in MAFFT was selected to generate pairwise alignment) (Katoh and Standley 2013; Muhire *et al*. 2014). The sequences used for the SDT analysis started from the conserved_up region and extended up to the stop codon of the coding sequence.

To examine the sequence diversity of *wtf* genes that exist as tandem pairs, we selected 44 genes from 22 tandem pairs. These genes were chosen to represent maximum diversity while minimizing redundancy. For any locus, we included at most one tandem pair from the REF lineage ancestry and one tandem pair from the NONREF lineage ancestry. To construct a neighbor-joining tree, we used MEGA X (version 10.0.5) (Kumar *et al*. 2018). The tree was rooted and visualized using FigTree v.1.4.3 (http://tree.bio.ed.ac.uk/software/figtree/).

### Analyzing recent evolutionary events affecting *wtf* genes

Inferring recent evolutionary events requires the grouping of sequences based on phylogenetic relationship. To establish the phylogenetic relationship between different isolates at a *wts* locus, we calculated pairwise SNP distances of unique sequences surrounding the locus using SNPs called from Illumina sequencing data. The pairwise SNP distance was defined as the number of SNP differences per kb in noncoding sequences (including inter-CDS intervals and introns) within a 20-kb window extending outward from the proximal boundary of the closest neighboring CDS. These SNP distances were then used as input for the hclust function of the R stats package, which generated a dendrogram. To divide the dendrogram into groups, we utilized the cutree function of the R stats package and specified a cut height that corresponds to a SNP distance cutoff of 2.5 SNPs/kb. For each *wts* locus, we combined the groupings based on the upstream and downstream sequences to create a final grouping (referred to as phylogroups), where members of a phylogroup met the SNP distance cutoff for both sequences.

If members within a phylogroup had different numbers of *wtf* genes, the phylogroup was designated as having undergone gene number changes. By inferring the ancestral gene number using outgroups, we determined whether there had been an increase or decrease in gene number within the phylogroup. For members with the same number of *wtf* genes, we compared their sequences and referred to different sequences as alleles. To identify candidate alleles that may have undergone ectopic gene conversion, we selected alleles that showed more than 5 SNP differences relative to the consensus sequence of the phylogroup within a region no longer than 200 bp (a density of SNP differences 10 times the SNP distance cutoff used for establishing phylogroups). The phylogroups containing candidate alleles were manually inspected to ensure that the phylogroup assignment was correct. We used outgroups to determine the ancestral alleles (recipient alleles) for gene conversion events. Next, the segments that had undergone gene conversion in the derived alleles (products of gene conversion) were used as queries to search against all *wtf* gene sequences for gene conversion donors using BLASTN. The BLASTN hits with >99% identity and >99% length coverage were designated as donors. In cases where more than one hit met the donor criteria, we gave preference to the hit on the same genome.

Alleles not affected by ectopic gene conversion were compared to identify substitutions and indels. Alleles affected by the same gene conversion event were also compared to identify substitutions and indels. For substitutions and indels that caused functional type changes of typical *wtf* genes, we identified the ancestral allele using outgroups.

Supplementary Table 3 lists the evolutionary events that we identified, including those that altered the number of *wtf* genes per locus and those that changed the functional types of typical *wtf* genes.

### Sequence alignment analysis of all typical *wtf* genes

To gain insights into the patterns of sequence divergence among *wtf* genes, we performed a global sequence alignment analysis of all typical *wtf* genes (genes other than *wts7*, *wts1112a*, *wts14*, and *wts15*), using sequences starting from the conserved_up region and extending up to the stop codon of the coding sequence. The 33-bp repeat in the third exon and the 21-bp repeat in the last exon, both of which can have varying numbers of repeat units, interfere with sequence alignment. To address this, we used the consensus sequences of the repeat units (Eickbush *et al*. 2019) as queries for iterative BLASTN searches. This allowed us to identify and remove the repeat sequences from the *wtf* gene sequences. We also excluded 25 *wtf* genes shorter than 1.2 kb because they had experienced significant segmental deletions. With the remaining 982 sequences, a sequence alignment was generated using PASTA v1.8.5 (Mirarab *et al*. 2015) and manually adjusted. Finally, we applied the Percentage Identity coloring scheme of Jalview (Waterhouse *et al*. 2009) to highlight conserved regions across all typical *wtf* genes.

### Sequence similarity network (SSN) analysis of the divergent segments in typical *wtf* genes

To analyze the divergence patterns of sequences of each divergent segment or sub-segment in an alignment-free manner, we performed sequence similarity network (SSN) analysis (Atkinson *et al*. 2009). Pairwise BLASTN searches were conducted for all sequences of a segment or sub-segment. Sequences with a BLASTN hit longer than 30 bp were considered similar and connected by edges in the SSN. For the D2 segment, which is less divergent than other segments, we used stricter megaBLASTN searches. The resulting networks were visualized using the prefuse force directed layout in Cytoscape version 3.8.0.

### Annotation of Tf transposons and solo LTRs

In the *S. pombe* reference genome, *wtf* genes are often found in close proximity to solo LTRs that originate from Tf transposons (Bowen *et al*. 2003). To annotate Tf transposons and solo LTRs that are adjacent to *wtf* genes in other genomes, we ran RepeatMasker (-cutoff 200) using an input repeat library composed of the sequences of full-length Tf1 and Tf2 transposons, as well as 48 solo LTRs from the reference genome.

### Demarcation of *wtf* gene boundaries

To determine the outer boundaries of *wtf* genes, we searched for inter-locus sequence similarities that extended beyond the previously annotated *wtf* gene sequences. To accomplish this, we collected for each *wtf* gene a sequence starting from the left end of the CDS of the left-flanking gene and ending at the right end of the CDS of the right-flanking gene. We then used the bedtools maskfasta tool to mask the previously annotated *wtf* gene sequences (from the conserved_up region to the stop codon), as well as Tf transposon sequences and solo LTR sequences, in these collected sequences. We replaced the masked nucleotides of each Tf transposon or solo LTR with a single N. Subsequently, we conducted pairwise BLASTN (-task blastn) analysis on all sequences, with the exception of the sequences containing *wts22* and *wts23* in JB900, two genes involved in an inversion. We only considered BLAST hits with e-values less than 0.01. The resulting BLAST hits were then aligned using MAFFT and visualized using Jalview.

## Results and discussion

### Obtaining full repertoires of *wtf* genes in 32 *S. pombe* isolates

Due to the repetitive nature of *wtf* genes, regular short-read sequencing is not suitable for investigating their sequence diversity. To comprehensively survey the intraspecific diversity of *wtf* genes, we generated long-read sequencing data for 20 distinct *S. pombe* isolates and performed de novo genome assemblies. Earlier versions of these assemblies were previously utilized to analyze retrotransposon diversity (collaboration with the Wolf group) (Tusso *et al*. 2022). For this study, we generated updated versions of our isolate assemblies (Supplementary Fig. 1A-E). Additionally, we obtained long-read-based *S. pombe* genome assemblies from a previous study that encompassed 11 additional *S. pombe* isolates (Tusso *et al*. 2019). The sequences of *wtf* genes were extracted from the genome assemblies of the 31 isolates. We also included the published sequences of *wtf* genes of one additional isolate (JB916/ FY29033) in our analysis (Eickbush *et al*. 2019). Overall, we obtained sequences of all *wtf* genes except one in 32 distinct *S. pombe* isolates (Fig. 1A). Among these 32 isolates, 5 belong to the REF lineage (also known as the *Sp* lineage) which includes the reference genome isolate, 10 belong to the NONREF lineage (also known as the *Sk* lineage), and the remaining 17 isolates are mosaic strains resulting from the admixture between the REF and NONREF lineages (Supplementary Fig. 1F) (Tao *et al*. 2019; Tusso *et al*. 2019).

**Figure 1.**
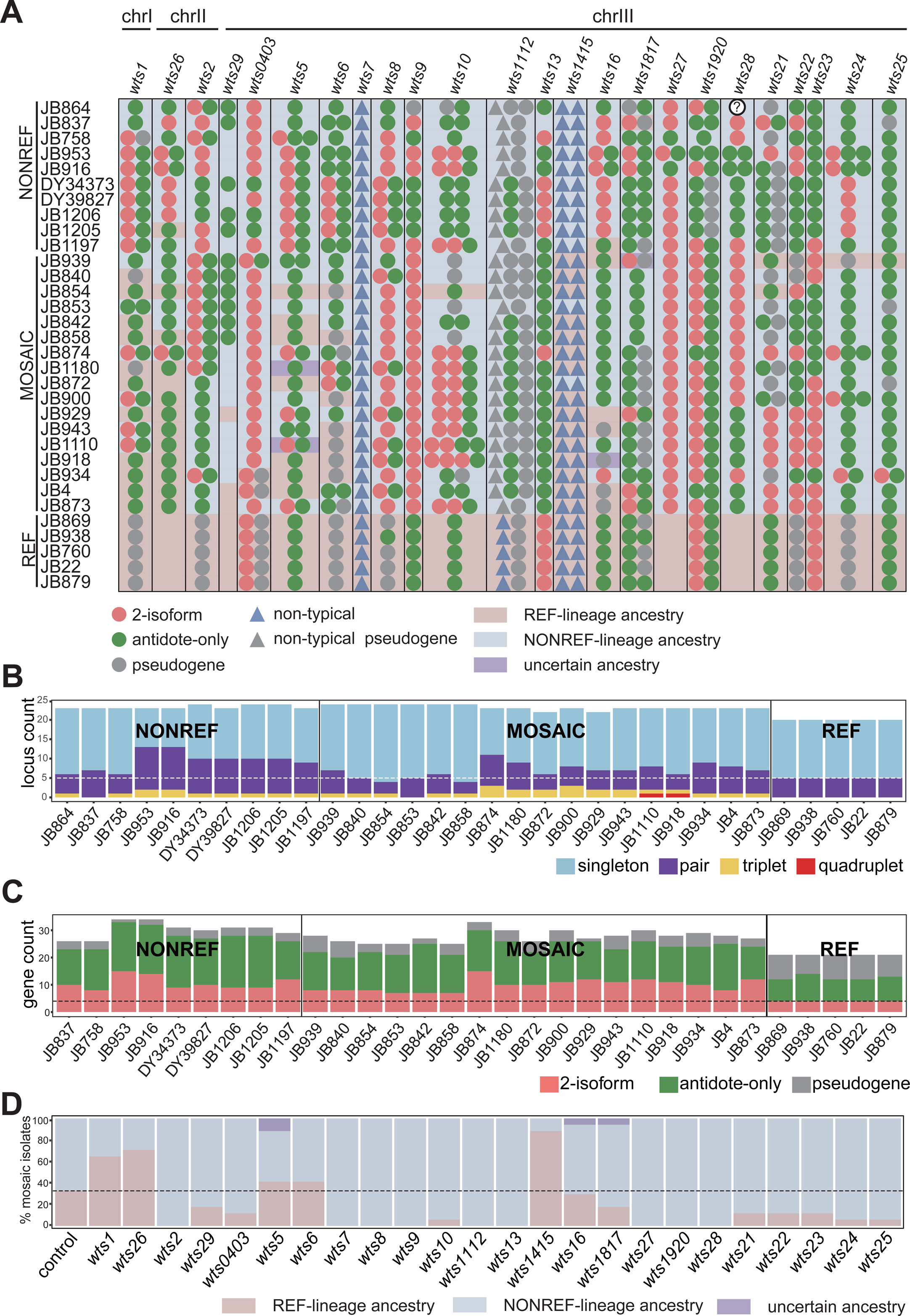
Identification and classification of *wtf* genes in 32 *S. pombe* isolates. (A) Overview of *wtf* genes in 32 isolates. Each row represents one isolate, and each column between two vertical black lines represents one *wts* locus. The order of *wts* loci corresponds to their positions in the reference genome. Color-filled circles indicate typical *wtf* genes (red for 2-isoform genes, green for antidote-only genes, and grey for pseudogenes). Color-filled triangles indicate non-typical *wtf* genes (blue for normal genes and grey for pseudogenes). In isolate JB864, a typical *wtf* gene is found at the *wts28* locus, but its exact sequence is uncertain. Therefore, it is represented by a white-filled circle with a question mark. The background color indicates the inferred lineage ancestry based on flanking SNPs. (B) Counts of the 4 types of *wtf* gene-containing loci in 32 isolates. Based on the number of *wtf* genes in a locus, *wtf* gene-containing loci are classified into 4 types: singleton, pair, triplet, and quadruplet. Each stacked bar represents the counts of the 4 types of loci for one isolate. The horizontal dashed line represents the counts of multi-gene loci in the REF lineage isolates. (C) Counts of the 3 types of typical *wtf* genes in 31 isolates. Typical *wtf* genes are classified into 3 types: 2-isoform, antidote-only, and pseudogene. Each stacked bar represents the counts of the 3 types of typical genes for one isolate. JB864 is not included because the exact sequence of the *wtf* gene at the *wts28* locus in JB864 is uncertain. The horizontal dashed line represents the counts of 2-isoform genes in the REF lineage isolates. (D) Percentages of mosaic isolates having REF-lineage ancestry or NONREF-lineage ancestry at each of the 24 *wts* loci. The leftmost bar shows the control percentages calculated using 100 randomly chosen 20-kb genomic regions. The horizontal dashed line represents the percentage of REF-lineage regions in this control calculation.

After obtaining the sequences of the *wtf* genes in the 32 *S. pombe* isolates, we proceeded to annotate their gene structures. To do this, we relied on the sequence conservation of these genes to the *wtf* genes in the reference genome, whose gene structures were previously annotated using a published long-read transcriptome dataset (Kuang *et al*. 2017; Eickbush *et al*. 2019). We classified genes that showed high similarity to *wtf7*, *wtf11*, *wtf14*, or *wtf15* in the reference genome as non-typical *wtf* genes. The remaining genes were classified as typical *wtf* genes. With the help of the gene structure annotations, we further divided the typical *wtf* genes into three categories: 2-isoform genes, antidote-only genes, and pseudogenes (Fig. 1A). All 2-isoform genes exhibit a hallmark feature—a conserved 150-bp sequence within intron 1 (Hu *et al*. 2017). This feature is always accompanied by an in-frame ATG codon situated either at the end of intron 1 or at the beginning of exon 2. This ATG codon is the start codon of the toxin isoform.

Two previous studies have characterized the complete sets of *wtf* genes in 4 *S. pombe* isolates, identifying a total of 24 genomic loci that harbor *wtf* genes, including 20 loci containing *wtf* genes in the reference genome isolate (Hu *et al*. 2017; Eickbush *et al*. 2019). Despite the inclusion of a substantially larger number of *S. pombe* isolates in this study, which more comprehensively represent the genetic diversity of *S. pombe*, no additional *wtf* gene-containing loci were discovered (Fig. 1A). This suggests that the 24 loci encompass all the genomic loci that host *wtf* genes in *S. pombe*. Based on this finding, we have designed a new naming scheme for *wtf* genes to reflect the loci they are associated with, allowing for consistent naming across isolates.

In this naming scheme, we first assign names to the loci containing *wtf* genes (Fig. 1A). Each locus is given a name with the prefix “*wts*” (for *wtf* site), followed by a number that corresponds to the number(s) in the name(s) of the *wtf* gene(s) present at that locus in the reference genome. For instance, the locus containing the *wtf1* gene in the reference genome is named *wts1*, while the locus hosting the tandem gene pair *wtf4* and *wtf3* (with *wtf4* upstream of *wtf3*) in the reference genome is named *wts0403* (the inclusion of two zeros ensures consistency with the names of the other loci harboring *wtf* gene pairs in the reference genome, which are *wts1112*, *wts1415*, *wts1817*, and *wts1920*). As for the 4 loci lacking *wtf* genes in the reference genome, we named them *wts26*, *wts27*, *wts28*, and *wts29*, in line with a naming scheme previously used for *wtf* genes in JB4/CBS5557 (Hu *et al*. 2017).

In our newly designed nomenclature, if a *wts* locus contains a single *wtf* gene, the name of the gene is the same as the name of the locus. If a *wts* locus contains multiple *wtf* genes oriented in tandem, the genes at that locus are named by combining the locus name with a suffix (*a*, *b*, *c*, or *d*) that indicates the relative positions of the genes. The suffix is assigned based on the order of the genes, going from the most upstream gene to the most downstream gene. Therefore, according to this nomenclature, *wtf4* and *wtf3* in the reference genome are referred to as *wts0403a* and *wts0403b*, respectively.

### The REF lineage has lower fraction of 2-isoform *wtf* genes and higher fraction of *wtf* pseudogenes than the NONREF lineage

The number of *wts* loci that contain *wtf* genes varies among isolates, ranging from 20 in all 5 REF-lineage isolates to 24 in some NONREF-lineage isolates and mosaic isolates (Fig. 1B). The number of *wtf* genes in an isolate also varies, ranging from 25 in all 5 REF-lineage isolates to 38 in 2 NONREF-lineage isolates (JB916 and JB953) (Fig. 1C). This larger variation in gene number is primarily due to differences in the number of *wts* loci that harbor multiple *wtf* genes (hereafter referred to as multi-gene loci). All REF-lineage isolates have 5 multi-gene loci, whereas all NONREF-lineage isolates have more than 5 multi-gene loci, with a maximum of 13. Among the 17 mosaic isolates, 13 have more than 5 multi-gene loci, with a maximum of 11 (Fig. 1B). Furthermore, while all multi-gene loci in the REF-lineage isolates contain gene pairs, NONREF-lineage isolates and mosaic isolates have multi-gene loci that contain gene triplets and even gene quadruplets (Figs. 1A and 1B).

The number of 2-isoform genes per isolate varies greatly, ranging from as few as 4 in 5 REF-lineage isolates to as many as 15 in the mosaic strain JB874 and a NONREF-lineage isolate JB953 (Fig. 1C). In terms of the proportion of typical genes, REF-lineage isolates have a lower percentage of 2-isoform genes (19%) compared to NONREF-lineage isolates (29%–44%) and mosaic isolates (26%–45%) (Fig. 1C). Additionally, REF-lineage isolates have a higher percentage of pseudogenes among typical genes (33%–43%) than NONREF-lineage isolates (3%–12%) and mosaic isolates (4%–23%) (Fig. 1C). It is known that the REF lineage has much lower intra-lineage diversity than the NONREF lineage, possibly due to a severe bottleneck the REF lineage went through during the most recent ice age (Tao *et al*. 2019; Tusso *et al*. 2019). The observed characteristics of the *wtf* genes in the REF lineage may be attributed to a founder effect or genetic drift, both of which are more pronounced in small populations.

Mosaic isolates result from the admixture between the REF lineage and the NONREF lineage. During the admixture, the REF-lineage allele and the NONREF-lineage allele of a *wts* locus may compete to be inherited by the admixed progeny, causing a biased inheritance ratio. To investigate this, we classified each *wts* locus in an isolate as having either REF-lineage ancestry or NONREF-lineage ancestry based on the SNPs in the unique sequences flanking the locus (Fig. 1A). We then calculated the percentages of mosaic isolates with REF-lineage ancestry or NONREF-lineage ancestry for each *wts* locus (Fig. 1D). As a control, we calculated the average percentage of mosaic isolates with REF-lineage ancestry in 100 randomly chosen 20-kb genomic regions, and this percentage was found to be 31%. Remarkably, for 9 out of the 24 *wts* loci, 0% (0/17) of mosaic isolates have REF-lineage ancestry. For more than half of the remaining 15 *wts* loci, the percentage of mosaic isolates with REF-lineage ancestry is notably lower than the control percentage. These results suggest that NONREF-lineage alleles tend to be preferentially inherited during inter-lineage admixture. Possible reasons for this preference include: a higher fraction of *wts* loci lacking *wtf* genes in the REF lineage, a higher fraction of loci containing only *wtf* pseudogenes in the REF lineage, a lower fraction of loci containing 2-isoform genes in the REF lineage, and/or possibly lower transmission distorting activity of REF-lineage 2-isoform genes.

### Sequence diversity of *wtf* genes

To analyze the diversity of *wtf* genes in the 32 *S. pombe* isolates, we calculated pairwise nucleotide identities of syntenic *wtf* genes in different isolates (Supplementary Fig. 3A). These inter-isolate comparisons were limited to sequences in syntenic loci with the same number of *wtf* genes. It is known that the genome-wide inter-isolate nucleotide identities among REF-lineage isolates, among NONREF-lineage isolates, and between REF-lineage and NONREF-lineage isolates are approximately 99.9%, 99.4%, and 99.0%, respectively (Jia *et al*. 2023). We observed that nucleotide identities of *wtf* genes in inter-isolate comparisons are often much lower than the genome-wide nucleotide identities, sometimes as low as around 60% (Supplementary Fig. 3A). This substantial divergence of syntenic *wtf* genes aligns with the model that *wtf* genes undergo dramatic sequence changes through non-allelic gene conversion (Hu *et al*. 2017; Eickbush *et al*. 2019). As expected, comparisons among the low diversity REF-lineage isolates (REF-REF comparisons) generally showed higher nucleotide identities of *wtf* genes than comparisons among the high diversity NONREF-lineage isolates (NONREF-NONREF comparisons). Comparisons across the two lineages (REF-NONREF comparisons) tended to have the lowest identities (Supplementary Fig. 3A). Notably, for many loci, the distribution of identity values indicates the existence of a limited number of distinct alleles.

We noticed that in 3 loci (*wts29*, *wts7*, *wts1415*), only high inter-isolate nucleotide identities are observed (Supplementary Fig. 3A). For the *wts7* and *wts1415* loci, which are the only two loci harboring exclusively non-typical *wtf* genes, the high identities are presumably due to the dissimilarity of non-typical *wtf* genes to other *wtf* genes preventing non-allelic gene conversion. For the *wts29* locus, the reason may be its special location. The *wts29* locus is located between *SPCC1884.01* and *nic1*, less than 20 kb away from the rDNA array at the left end of chromosome III. This proximity to a recombination-repressed rDNA region may hinder non-allelic gene conversion.

We also calculated pairwise nucleotide identities of typical *wtf* genes within the same isolate (intra-isolate comparisons) (Supplementary Fig. 3B). For all isolates, the identity values show a broad distribution ranging from just above 50% to 100%, with the median falling at around 75%. We paid special attention to multi-gene loci. If multi-gene loci result from local duplication of *wtf* genes, we would expect the identities within the same multi-gene locus (intra-locus identities, including within-pair, within-triplet, and within-quadruplet comparisons) to be higher than those between different loci (inter-locus identities). However, contrary to this expectation, we observed that intra-locus identities tend to be lower than inter-locus identities (Supplementary Fig. 3B). This suggests that multi-gene loci, in general, do not originate from local gene duplication.

To further investigate multi-gene loci, we constructed a phylogenetic tree of 44 *wtf* genes from 22 representative *wtf* gene pairs (Supplementary Fig. 3C). In this tree, the two *wtf* genes from each gene pair are consistently found far apart, which matches the low within-pair identities shown in Supplementary Fig. 3B. Notably, the upstream *wtf* genes in these gene pairs (with 2 exceptions) cluster together in the tree. Similarly, the downstream *wtf* genes in the gene pairs (with 1 exception) also cluster together. Remarkably, all 2-isoform genes in these gene pairs are upstream genes. These patterns suggest that the upstream and downstream genes may have distinct origins. It is conceivable that all gene pairs descend from an ancestral gene pair, in which the upstream gene and the downstream gene are distinct from each other.

### Inferring recent evolutionary events altering *wtf* genes

Previous studies on the evolution of *S. pombe wtf* genes did not attempt to determine the exact evolutionary events that occurred to *wtf* genes (Hu *et al*. 2017; Eickbush *et al*. 2019). In this study, we developed a strategy to infer recent evolutionary events that have altered *wtf* genes using our large dataset of *wtf* gene sequences. First, for each *wts* locus, we established the phylogenetic relationship of different isolates using pairwise SNP divergence in flanking genomic regions. We then categorized the isolates into phylogroups based on a divergence cutoff of 2.5 SNPs per kb. This cutoff was chosen to allow for variations in *wtf* genes within a phylogroup while minimizing situations where more than 2 alleles exist within a phylogroup. When applying this cutoff, REF-lineage isolates consistently fell into the same phylogroup, while NONREF-lineage isolates typically fell into multiple phylogroups. Next, we compared the alleles within a phylogroup to the alleles in outgroups. The allele identical or closely related to allele(s) in outgroups was classified as the ancestral allele, and the allele sharing weaker similarities with outgroups was classified as the derived allele resulting from a recent evolutionary event. For this analysis and the analyses described later, we supplemented the dataset with 131 additional *wtf* gene sequences obtained by amplicon sequencing (Supplementary Table 2). These sequences were from *S. pombe* isolates distinct from the 32 isolates described above.

We identified a total of 9 recent evolutionary events that altered the number of *wtf* genes per locus (Fig. 2A and Supplementary Table 3). Among these events, 3 led to an increase in gene number, while 6 resulted in a decrease in gene number. The 6 events that decreased gene number can all be explained by recombination between two adjacent genes within a multi-gene locus (Supplementary Fig. 4A). The 3 events that increased gene number may have occurred in two different ways: ectopic insertion and unequal crossing over. At the *wts0403* locus, the ancestral state of the phylogroup that includes JB939 is a single *wtf* gene. However, JB939 has two genes at this locus, *wts0403a* and *wts0403b*. Sequence alignment revealed that the *wts0403b* gene is the newly added gene (Supplementary Fig. 4B). BLASTN analysis indicated that the 5′ portion and the 3′ portion of *wts0403b* perfectly match *wts5b* and *wts5a* in JB939, respectively. One possible model explaining the origin of *wts0403b* in JB939 is that *wts5a* and *wts5b* recombined and formed a circular DNA, which then ectopically inserted into the *wts0403* locus by recombination at a site downstream of the single gene originally present at this locus (Supplementary Fig. 4C). The other two gene number-increasing events both occurred at the *wts10* locus and resulted in the formation of a gene quadruplet. The quadruplet in JB1110 likely arose from unequal crossing over between a gene pair and a gene triplet, two alleles present in the same phylogroup (Supplementary Figs. 4B and 4D). The quadruplet in JB918, belonging to a different phylogroup, was probably generated through unequal crossing over between two copies of a gene triplet (Supplementary Figs. 4B and 4E).

**Figure 2.**
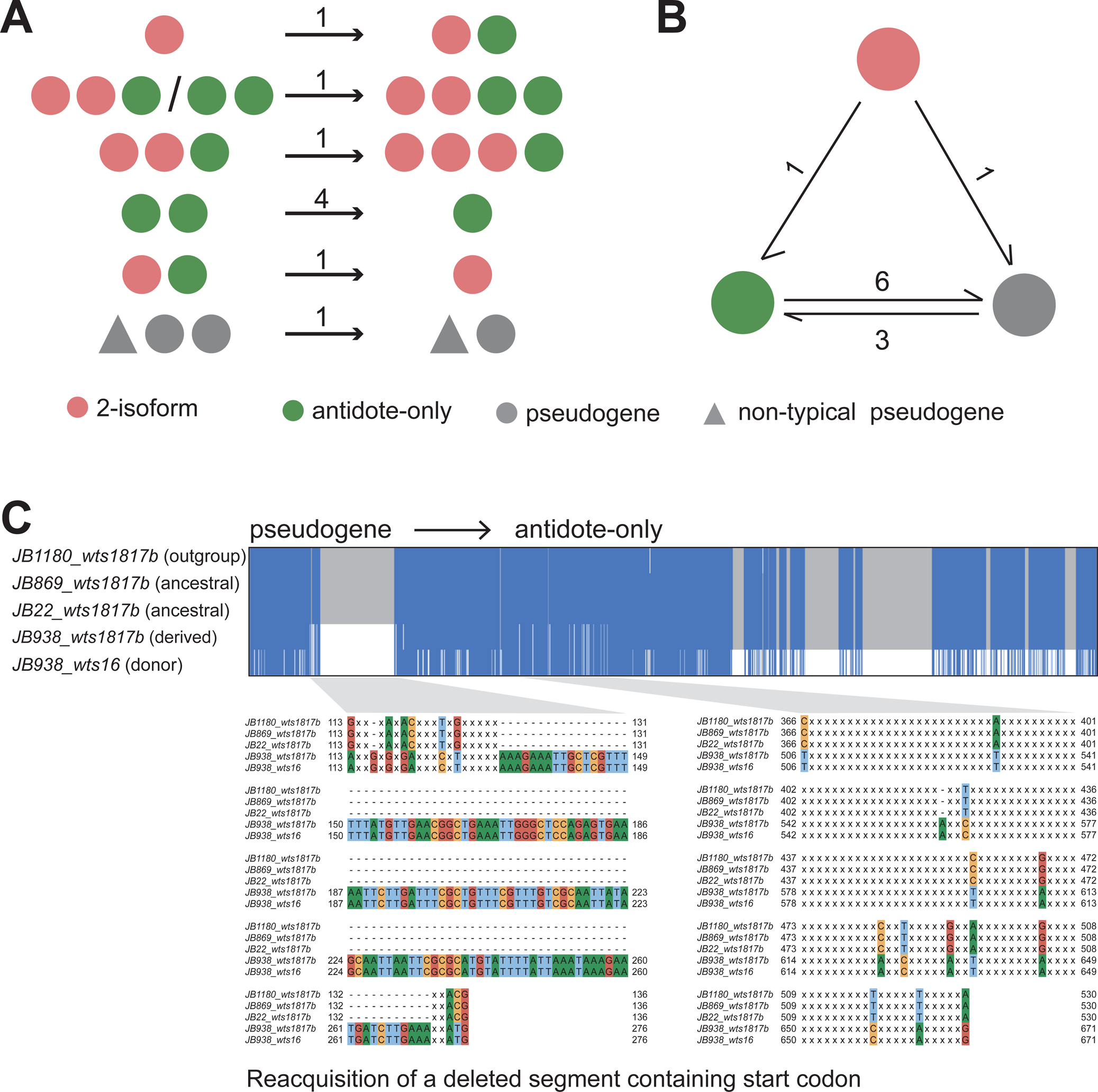
Recent evolutionary events altering the number of *wtf* genes per locus and changing the functional type of *wtf* genes. (A) Schematic depicting the functional types of the genes in the ancestral allele(s) and the derived allele for the 11 events altering the number of *wtf* genes per locus. The arrows point from the ancestral allele(s) to the derived allele. The number on top of each arrow indicates the number of events with the depicted pattern. Details of these gene number-altering events are shown in Supplementary Fig. 4. (B) Schematic depicting the functional types of the genes in the ancestral allele(s) and the derived allele for the 9 events altering the functional type of *wtf* genes. Except for 3 substitution/indel events that convert antidote-only genes to pseudogenes and 1 indel event that converts a two-isoform gene to a pseudogene, the remaining 5 events are non-allelic gene conversion events, including 3 events that convert pseudogenes to antidote-only genes, 1 event that converts an antidote-only gene to a pseudogene, and 1 event that converts a two-isoform gene to an antidote-only gene. Details of the 5 gene conversion events are shown in Supplementary Fig. 5. (C) The revival of a pseudogene through non-allelic gene conversion. Top, a Jalview-generated alignment overview illustrating where gene conversion occurred. Two regions underwent gene conversion. In the region on the left, the ancestral allele contains a deletion, while the derived allele no longer contains the deletion. Bottom, zoomed-out views of the alignment showing the nucleotide sequences in the regions that underwent gene conversion. In these zoomed-out views, identical nucleotides are represented as “x”.

For recent evolutionary events that did not alter the number of genes in a locus but altered the sequence of a *wtf* gene, we focused on the events that resulted in a change of functional type for a typical *wtf* gene, based on the classification of typical *wtf* genes into three functional types: 2-isoform, antidote-only, and pseudogene. We identified 11 such events (Fig. 2B and Supplementary Table 3). We found 2 substitution events and 4 indel events that resulted in the transformation of 5 antidote-only genes and a 2-isoform gene into pseudogenes (Supplementary Table 3). Furthermore, we discovered a gene conversion event that caused a 2-isoform gene to become an antidote-only gene, and another gene conversion event that caused an antidote-only gene to become a pseudogene (Supplementary Fig. 5 and Supplementary Table 3). Interestingly, we also identified 3 gene conversion events that transformed pseudogenes into antidote-only genes (Figure 2C, Supplementary Fig. 5, and Supplementary Table 3). These findings indicate that the revival of dead *wtf* genes through gene conversion can occur and may have played a crucial role in the persistence of *wtf* genes during evolution.

### Divergent segments in *wtf* genes originated prior to the REF-NONREF lineage split

A previous study reported that phylogenetic analyses of individual exons of typical *wtf* genes always yielded a tree with two main clades, but the grouping of genes into these main clades was not consistent across different exons (Eickbush *et al*. 2019). This suggests that typical *wtf* genes contain multiple segments that exhibit a two-clade divergence pattern. It is unclear, though, whether these segments strictly correspond to exons. To investigate this, we generated an alignment of the nucleotide sequences of 930 typical *wtf* genes, encompassing nearly all typical *wtf* genes presented in the 32 isolates and the amplicons (Fig. 3A). The alignment starts from the beginning of the conserved_up region, an approximately 288-bp highly conserved sequence upstream of exon 1 (Hu *et al*. 2017), and ends at the stop codon.

**Figure 3.**
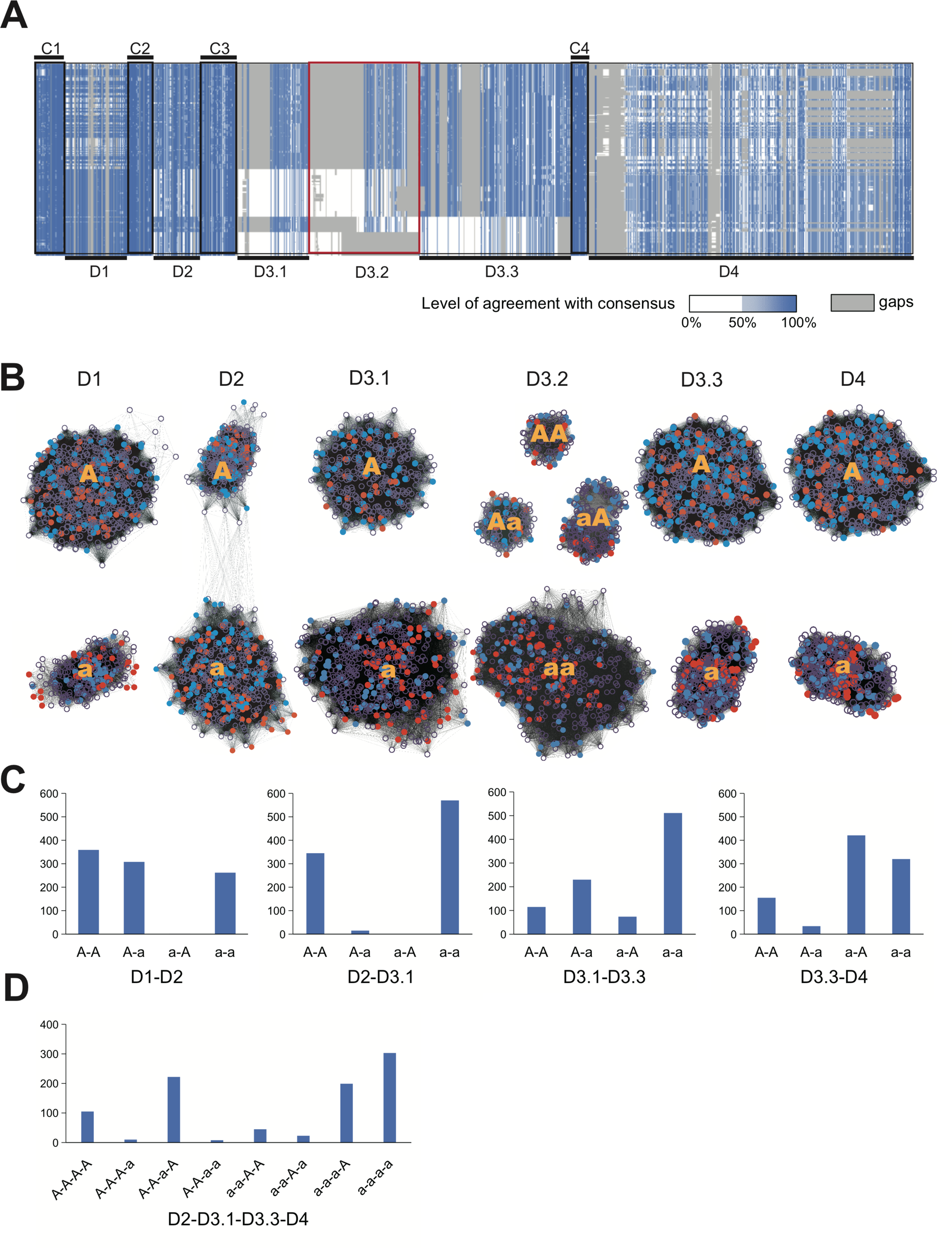
Divergent segments in *wtf* genes originated before the REF-NONREF split. (A) Nucleotide sequence alignment of typical *wtf* genes. The alignment is shown as a Jalview-generated overview. Gaps in the alignment are shown in dark grey. Nucleotides are colored according to the percentage agreement with the consensus. (B) Sequence similarity network (SSN) analysis of divergent segments and subsegments of *wtf* genes. Each node represents one *wtf* gene. Lines between nodes indicate sequence similarity detected by BLAST. Nodes representing *wtf* genes from 5 REF-lineage isolates (JB22, JB760, JB869, JB879, and JB938) are colored red. Nodes representing *wtf* genes from 5 NONREF-lineage isolates (JB758, JB864, JB916, JB953, and JB1206) are colored blue. Nodes representing *wtf* genes from other isolates are shown as empty circles. (C) The number of combinations of sequence types of adjacent divergent segments/ sub-segments that have 2 sequence types (denoted as “A” and “a”, respectively). (D) The number of combinations of sequence types of D2, D3.1, D3.3, and D4. The combinations between D2 and D3.1 are limited to the two predominant ones (A-A and a-a).

The alignment revealed 4 short regions showing strong nucleotide sequence conservation across all typical *wtf* genes. These regions, referred to as “conserved segments”, were named C1-C4. C1 corresponds to the most 5′ portion of the conserved_up region, C2 corresponds to a part of the conserve_up region near its 3′ end, C3 includes the 3′ portion of exon 1 and the exon 1-intron 1 junction, and C4 spans the junction of intron 2 and exon 3. The regions located between these conserved segments and the region downstream of C4 exhibited strong divergence patterns. These regions, referred to as “divergent segments”, were named D1-D4. D1 is located within the conserved_up region, D2 starts from the most 3′ portion of the conserved_up region and ends within exon 1, D3 includes intron 1, exon 2, and part of intron 2, and D4 starts from within exon 3 and extends to the stop codon. In the alignment, D1, D2, and D4 each exhibited two highly divergent sequence types, while D3 showed a more complex diversity pattern. We further divided D3 into 3 sub-segments: D3.1 (the 5′ portion of intron 1), D3.2 (the 3′ portion of intron 1 and the 5′ portion of exon 2), and D3.3 (the 3′ portion of exon 2 and the 5′ portion of intron 3). D3.1 and D3.3 each showed 2 distinct sequence types, whereas D3.2 exhibited 4 different sequence types. Interestingly, the 4 sequence types of D3.2 strictly correlated with the 4 combinations of the sequence types of D3.1 and D3.3. The divergent segments in typical *wtf* genes likely account for the previously observed two-clade pattern when analyzing individual exons.

To validate the findings obtained from sequence alignment, we utilized an alignment-free approach known as sequence similarity network (SSN) (Atkinson *et al*. 2009). In an SSN, sequences are represented as nodes, and BLAST-identified pairwise similarities are depicted as edges connecting the nodes. We employed this method to analyze the nucleotide sequences of divergent segments/sub-segments of typical *wtf* genes (Fig. 3B). The results demonstrated that, in line with the divergence patterns observed in the sequence alignment, the sequences of D1, D2, D3.1, D3.3, and D4 formed 2 distinct clusters, whereas the D3.2 sequences formed 4 clusters.

When did the two clades of divergent segments/sub-segments arise in evolution? Since all currently known *S. pombe* isolates descend from two ancient lineages—the REF lineage and the NONREF lineage—we hypothesized that the two clades of divergent segments/sub-segments may share a common origin and diverged from each other after the split of these two lineages. To test this possibility, we highlighted the sequences from 5 REF lineage isolates and 5 NONREF lineage isolates in the SSN (Fig. 3B). We found that for all divergent segments/sub-segments, sequences from both lineages co-exist in each cluster in the SSN, thus rejecting our hypothesis. The two clades of divergent segments/sub-segments most likely originated before the REF-NONREF lineage split.

From inspecting the sequence alignment shown in Fig. 3A, we noticed that the sequence types in different divergent segments/sub-segments are not randomly mixed. For instance, it appears that each of the 2 sequence types in D2 is mostly associated with only one of the sequence types in D3.1. To facilitate the investigation of this non-random association, we chose to use the letters “A” and “a” to distinguish the two sequence types of D1, D2, D3.1, D3.3, and D4. We arbitrarily assigned the letter “A” to the sequence type of D1, D2, D3.1, D3.3, and D4 in the gene *wtf13* in the reference genome, which is an active driver gene (Bravo Núñez *et al*. 2018a) (Fig. 3B). The sequence types of D3.2 are denoted by “AA”, “Aa”, “aA”, and “aa” to reflect the fact that these 4 sequence types of D3.2 strictly correlate with the 4 combinations of the sequence types of D3.1 and D3.3.

We counted the number of different combinations of sequence types of adjacent divergent segments/sub-segments that have 2 sequence types (Fig. 3C). Consistent with the visual impression of the alignment, almost all combinations between D2 and D3.1 are either A-A or a-a combinations. Interestingly, there is no a-A combination among the combinations between D1 and D2, despite the other 3 combinations being found at similar frequencies. Strikingly, the non-random association of sequence types is not confined to adjacent divergent segments/sub-segments. We discovered that the A-A combination between D2 and D3.1 is predominantly linked to sequence type A of D4 (Fig. 3D). The exact reasons for the non-random association of sequence types are unclear. We speculate that certain combinations may not be compatible with the proper functions of the protein products.

### LTRs shape the upstream sequences of *wtf* genes

*wtf* genes were so named because they are often found near the solo LTRs of Tf retrotransposons (Wood *et al*. 2002; Bowen *et al*. 2003). Bowen et al. suggested that the reason for this frequent association could be that an ancestral *wtf* gene and its flanking LTR(s) were duplicated together to new sites in the genome during the expansion of the *wtf* gene family. According to this proposal, the *wtf*-associated LTRs would be more closely related to each other than to LTRs not associated with *wtf* genes. However, a thorough phylogenetic analysis of the LTRs in the reference genome showed that this is not the case (Bowen *et al*. 2003). Despite the lack of concrete evidence supporting the possibility of the co-duplication of *wtf* genes and LTRs to new genomic locations, this idea has been further developed by Eickbush et al., who proposed that the number of *wtf* gene-containing loci may have increased through recombination-mediated duplication of LTR-flanked *wtf* genes to pre-existing LTRs elsewhere in the genome (Eickbush *et al*. 2019). This model predicts that sequences on the *wtf*-gene-proximal side of the LTRs should share strong similarities across different loci. We investigated whether this prediction is true.

We first applied RepeatMasker to annotate the LTRs in the long-read-based genome assemblies of *S. pombe* isolates (Smit *et al*. 2015). We then selected 7 representative isolates, including 1 REF-lineage isolate (JB22), 3 NONREF-lineage isolates (JB758, JB864, and JB953), and 3 mosaic isolates (JB4, JB872, and JB1180), to perform in-depth analysis of the sequences on the *wtf*-gene-proximal side of the LTRs. To avoid uncertainty in demarcating the boundaries of LTRs with highly degenerated ends, we only considered LTRs with an intact end on the side facing the adjacent *wtf* genes. The intact ends of LTRs can be easily recognized because the LTRs of Tf retrotransposons in *S. pombe* contain a trinucleotide inverted repeat (5′-TGT…ACA-3′) at their ends. As noted before by Bowen et al. (Bowen *et al*. 2003), LTRs located at the upstream side of *wts* loci are always in close proximity to the upstream end of the conserved_up region (Fig. 4A). We referred to the interval between an upstream LTR and the conserved_up region as BLC (between LTR and conserved_up). There are 2 *wts* loci (*wts1920* and *wts27*) that have a reverse-oriented upstream LTR. Interestingly, the BLCs at these 2 loci share largely the same 87-bp sequence (Fig. 4A).

**Figure 4.**
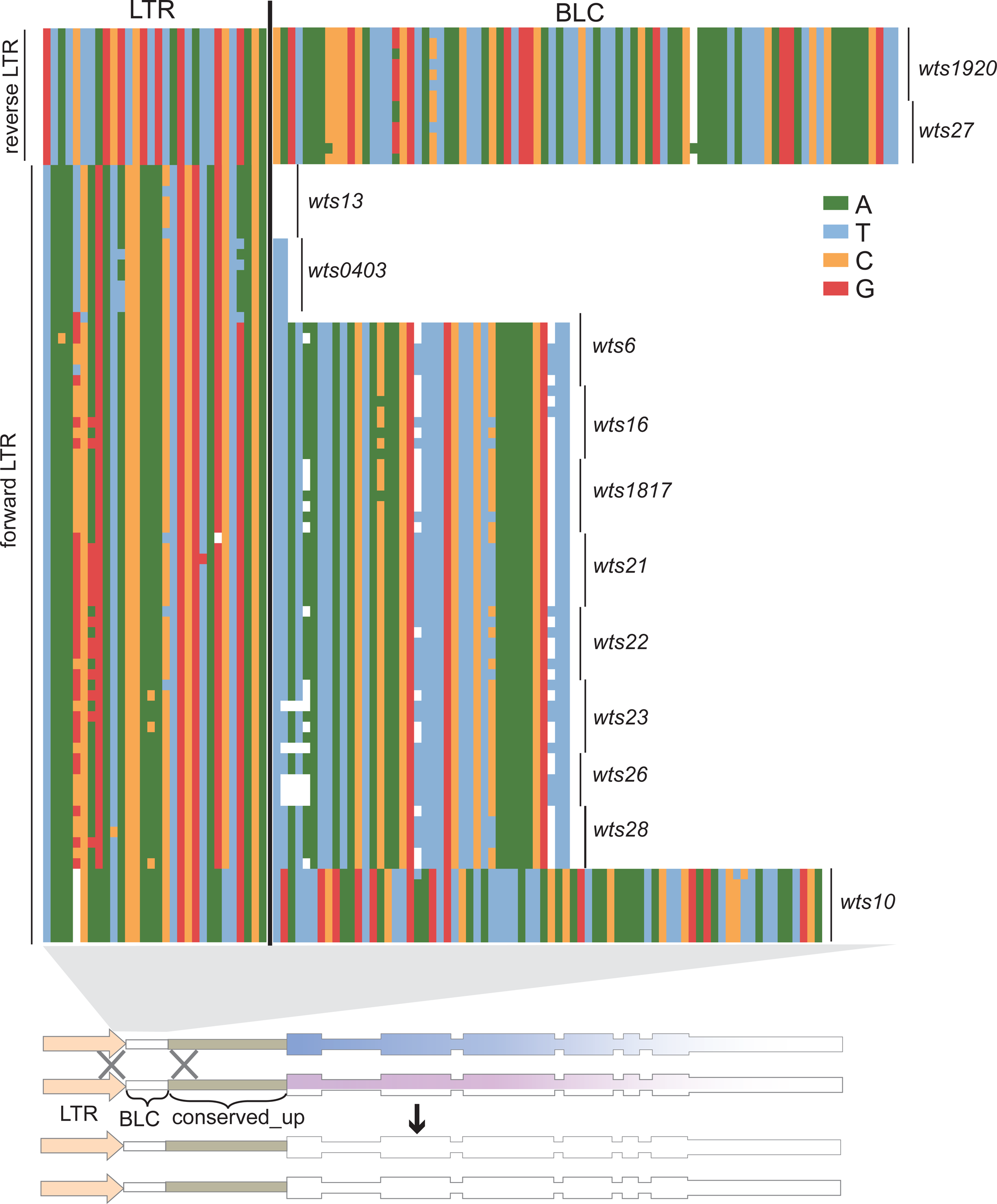
LTRs shape the upstream sequences of *wtf* genes. At the top, an alignment of nucleotide sequences upstream of the conserved_up region is presented. Each sequence includes a 30-bp *wtf*-gene-proximal portion of the LTR on the upstream side of a *wts* locus and the BLC sequence between the LTR and the conserved_up region. The sequences are from 7 representative isolates and are ordered from top to bottom as follows: JB22, JB1180, JB4, JB758, JB864, JB953, and JB872. For the *wts27* and *wts28* loci, there are only 6 sequences as JB22 does not harbor *wtf* genes at these two loci. For the *wts26* locus, there are only 5 sequences as JB22 and JB872 do not harbor *wtf* genes at this locus. At the bottom, a schematic shows how LTRs upstream of *wts* loci can act as homology arms to homogenize the upstream sequences of *wtf* genes through non-allelic gene conversion.

There are 11 *wts* loci with a forward-oriented upstream LTR (Fig. 4A). At the *wts13* locus, the upstream LTR is immediately adjacent to the conserved_up region and thus there is no BLC sequence. For the other 10 loci, there are 3 types of BLC sequences: a dinucleotide TT sequence at the *wts0403* locus in all isolates and at the *wts6* locus in JB22, a 39-bp sequence and its variants at 8 different loci (*wts6*, *wts16*, *wts1817*, *wts21*, *wts22*, *wts23*, *wts26*, and *wts28*), and a 74-bp sequence at the *wts10* locus. One possible explanation of the sharing of the same BLC sequence across different loci is that the BLC sequence may have originated in one locus and been transferred to new locations through recombination. This lends support to the model that *wtf* genes and adjacent LTRs have been co-duplicated to pre-existing LTRs.

However, when we examined the LTRs located downstream of *wtf* loci, we found a different situation (Supplementary Fig. 6). There are 3 loci (*wts1*, *wts0403*, and *wts27*) that have a reverse-oriented downstream LTR. We did not find any obvious sharing of identical sequence on the *wtf*-gene-proximal side of the LTRs between these 3 loci. Additionally, there are 4 loci (*wts9*, *wts13*, *wts22*, and *wts23*) that have a forward-oriented downstream LTR. Similarly, the sequences on the *wtf*-gene-proximal side of the LTRs at these loci are different. Therefore, we were unable to obtain definitive evidence supporting the model that *wts* loci as a whole have been duplicated to new locations through recombination with pre-existing LTRs at the new locations. Instead, we propose that LTRs upstream of *wts* loci can act as homology arms, homogenizing the upstream sequences of *wtf* genes through non-allelic gene conversion (Fig. 4B). The reason why LTRs have not caused the same homogenizing effect on sequences downstream of *wts* loci is unclear. It is worth noting that BLC sequences have been shown to be important for the full antidote activities of two *wtf* driver genes (Hu *et al*. 2017). It is plausible that selection has played a role in preserving the upstream sequence alterations resulting from LTR-mediated gene conversion.

### Identification of a type of direct repeats flanking *wtf* genes

Previous studies of *wtf* genes have mainly focused on sequences from the beginning of the conserved_up region to the stop codon. Consequently, it remains unclear where the outer boundaries of *wtf* genes lie. To address this, we conducted a BLASTN analysis to identify inter-locus similarities in the sequences upstream of the conserved_up region and downstream of the stop codons using BLASTN analysis. For the upstream sequences, there are hardly any obvious inter-locus similarities except for the short BLC sequences between an upstream LTR and the conserved_up region. Interestingly, our analysis revealed that downstream of the stop codons of typical *wtf* genes, there are sequences of several hundred base pairs that exhibit similarities across the majority of loci. We refer to these sequences as conserved_down sequences.

We generated an alignment of the conserved_down sequences (Supplementary Fig. 7). In this alignment, the sequences immediately downstream of the stop codons showed a divergent pattern consisting of two sequence types. These two sequence types correspond to the two clades of D4, which is the most 3′ divergent segment described earlier, starting from within exon 3 and ending at the stop codon. Thus, this region can be considered an extension of the D4 segment. Beyond this region, there is an approximately 310-bp sequence that exhibits strong similarity across *wts* loci of both D4 clades (Supplementary Fig. 8). We named this conserved region C5. It is worth noting that not all *wts* loci have downstream C5 regions, and among those that do, some have 3′ truncated C5. Retrotransposon-mediated deletions could potentially account for the loss and truncation of C5, as solo LTRs are frequently observed at the downstream boundaries of the *wts* loci wherein a complete downstream C5 is absent. For loci that contain a complete downstream C5, there are no obvious inter-locus sequence similarities beyond C5, suggesting that C5 may represent the most downstream sequence of *wtf* genes.

The last 4 nucleotides of C5 are predominantly TAAG. Interestingly, for C5 located between *wtf* genes within a multi-gene locus, these 4 nucleotides are also the first 4 nucleotides of the conserved_up region of the downstream *wtf* gene. The majority of conserved_up regions, including those at singleton loci and those belonging to the most 5′ gene at multi-gene loci, begin with TAAG. Thus, *wtf* genes with a complete downstream C5 region often have the same TAAG sequence at both ends, and within multi-gene loci, two adjacent genes are frequently connected head-to-tail with an overlap of TAAG. This observation prompted us to investigate whether this TAAG sequence might be part of a longer repeat sequence surrounding *wtf* genes.

At 3 *wts* loci (*wts1*, *wts5*, and *wts24*), we found sequences upstream of the TAAG in the conserved_up region of the most 5′ gene that match the sequence upstream of TAAG in C5 (Fig. 5, Supplementary Figs. 9 and 10). The longest such sequence was found at the *wts1* locus and is approximately 160-bp long. Based on this observation, we propose that direct repeats of 160 bp, and possibly longer, may have flanked many ancestral *wtf* genes. We refer to these repeats as “outer repeats”. With the exception of the ones located between *wtf* genes within multi-gene loci, the outer repeats have eroded during evolution to the point that only 3 loci still contain obvious recognizable outer repeat sequences at their 5′ side. No outer repeats are present outside of *wts* loci. We speculate that the outer repeats may have played a role in the early expansion of the *wtf* gene family. One possibility is that the outer repeats may correspond to a type of dispersed repeats present in the genome of the ancestor of *S. pombe* and may have served as acceptors for the ectopic duplication of an ancestral *wtf* gene flanked by outer repeats, similar to how dispersed 5S rDNA genes facilitate the expansion of *wtf* genes in *S. octosporus* and *S. osmophilus* (De Carvalho *et al*. 2022).

**Figure 5.**
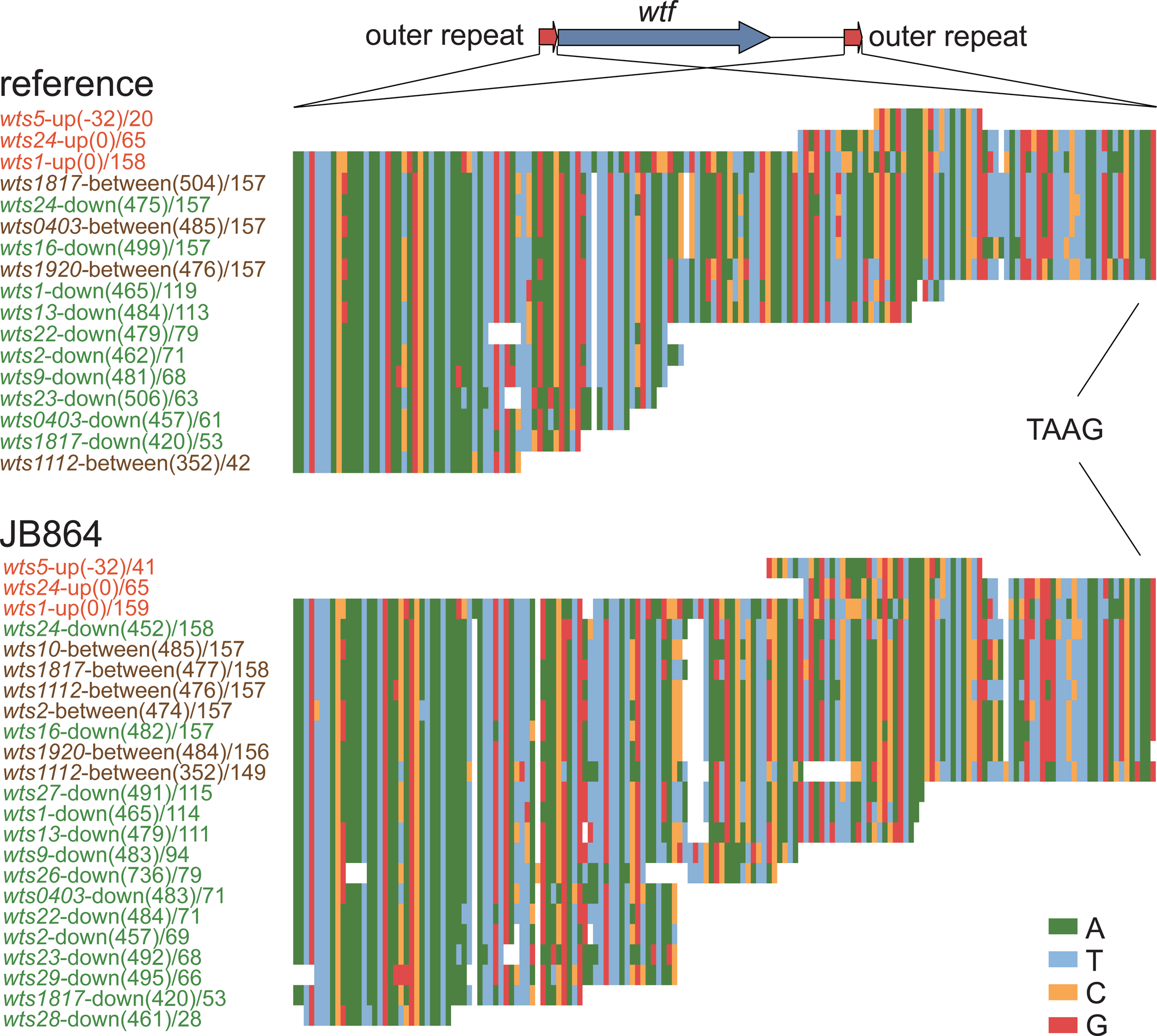
The outer repeats of *wtf* genes. Alignments of nucleotide sequences of the outer repeats in the reference genome and in the genome of JB864 are presented. The names of outer repeats on the upstream side of a *wts* locus contain the word “up” and are colored red. The names of outer repeats on the downstream side of a *wts* locus contain the word “down” and are colored green. The names of outer repeats located between two *wtf* genes within a multi-gene locus contain the word “between” and are colored brown. The numbers in brackets indicate the distances between the outer repeat and the conserved_up region (for upstream outer repeats) or the nearest upstream stop codon (for downstream outer repeats and outer repeats located between two *wtf* genes). Numbers after slashes indicate the lengths of the outer repeats. In the schematic at the top, two red arrows represent outer repeats and the blue arrow represents the *wtf* gene (from the beginning of the conserved_up region to the stop codon) located between the outer repeats.

## Conclusion

In summary, this study has provided a comprehensive analysis of the diversity of *wtf* genes in *S. pombe*, revealing a complex picture of their evolutionary dynamics and the influence of various factors on their sequence variation. Our analysis highlights lineage-specific differences in *wtf* gene composition, with the REF lineage exhibiting fewer 2-isoform genes and more pseudogenes, possibly due to historical population bottlenecks. Taking advantage of our large dataset, we have identified numerous recent evolutionary events that have altered the number and sequence of *wtf* genes. Notably, we observed that non-allelic gene conversion can revive pseudogenes, providing direct evidence of its role in the evolutionary persistence of this KMD family. Additionally, we have improved the understanding of the divergent segments within *wtf* genes and demonstrated that their divergence occurred prior to the REF-NONREF split. This study has also uncovered the role of LTRs in shaping the upstream sequences of *wtf* genes. Moreover, we have identified a new type of direct repeats surrounding *wtf* genes, which may have contributed to the expansion of *wtf* genes. Overall, this research offers new insights into the complex evolutionary dynamics of *wtf* genes in *S. pombe*.

## Acknowledgments

We thank Sergio Tusso and Jochen Wolf for sharing information and for discussions. We thank Jürg Bähler for making *S. pombe* natural isolate strains available through the Yeast Genetic Resource Center of Japan (YGRC/NBRP), and YGRC/NBRP for providing the strains. This work was supported by funding from the Ministry of Science and Technology of the People’s Republic of China, the Beijing municipal government, and Tsinghua University.

---

**Supplementary Figure 1.**
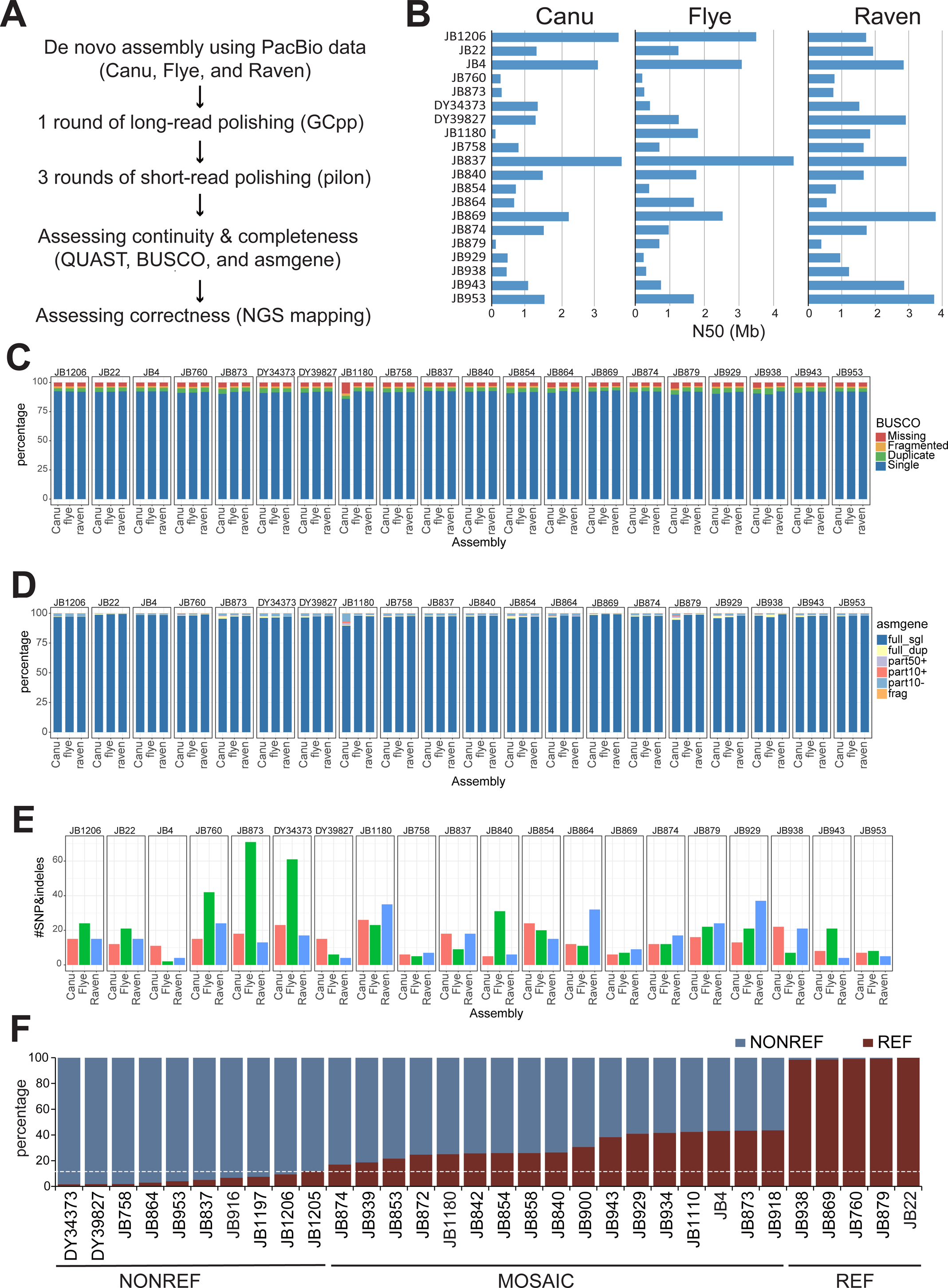
Quality assessment of long-read-based genome assemblies and the lineage ancestry of the 32 isolates. (A) Flowchart depicting the pipeline used in this study for de novo assembly of PacBio sequencing data of 20 *S. pombe* isolates and the subsequent assessment of assembly quality. (B) N50 values (an indicator of assembly contiguity) of the 60 assemblies reported by QUAST. (C) Assembly completeness assessment results reported by BUSCO. (D) Assembly completeness assessment results reported by asmgene. (E) Assembly correctness assessment by mapping Illumina sequencing reads to the assemblies and calling variants using GATK, Samtools, and DeepVariant. The barplot shows the number of SNPs and indels called by at least two variant calling tools. (F) Lineage ancestry of the 32 isolates whose *wtf* genes have been comprehensively analyzed in this study. The sequence of a representative assembly for each isolate was divided into 20-kb windows. The number of SNPs relative to JB22 (a REF lineage isolate) was calculated for each window. The percentage of REF lineage sequence in each isolate was determined by dividing the number of windows containing no more than 20 SNPs by the total number of windows. The dashed line denotes 10% REF lineage sequence, a threshold for distinguishing pure-lineage isolates and mosaic isolates (Tusso *et al*. 2019).

**Supplementary Figure 2.**
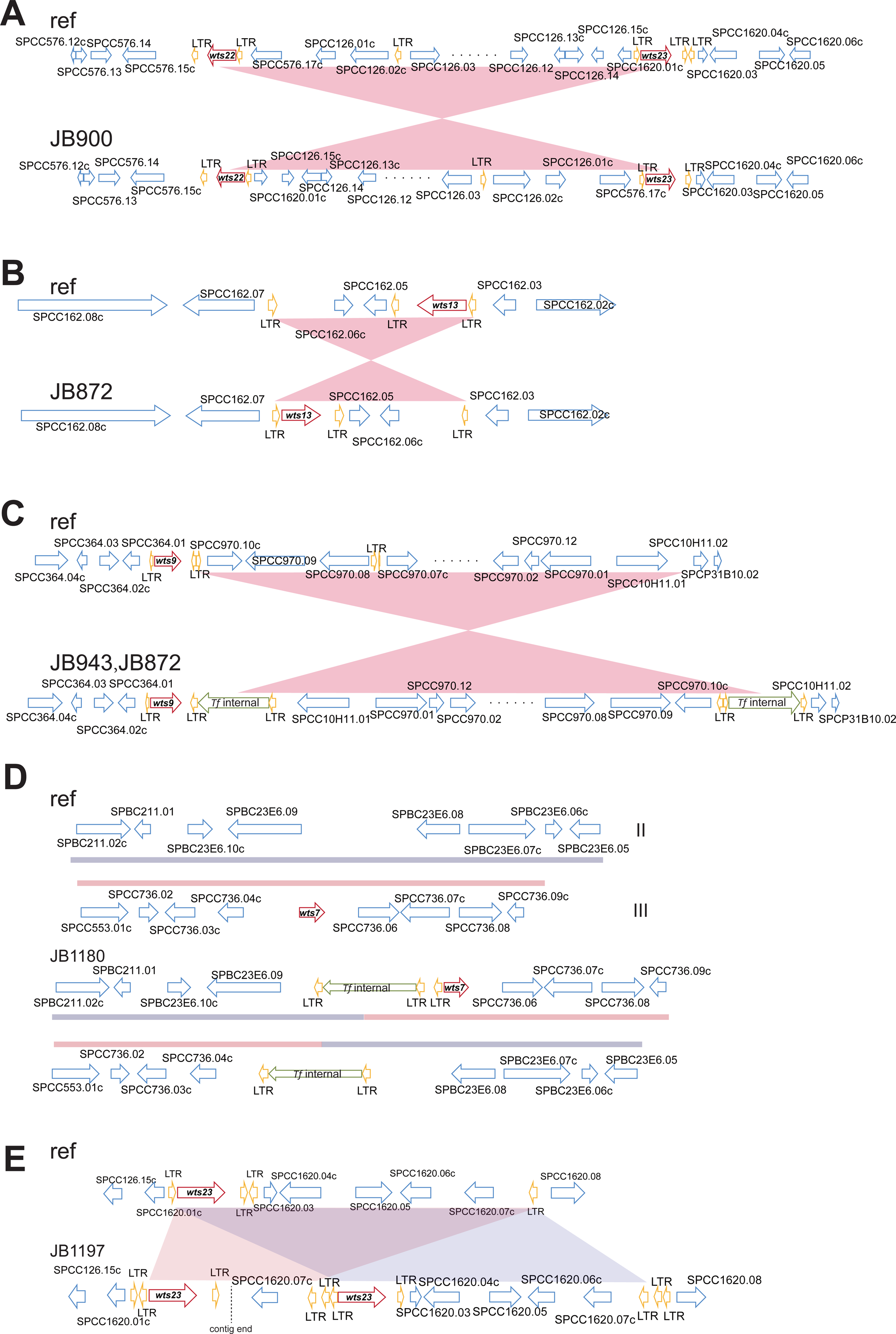
Schematics of genome rearrangements altering the order of unique genes surrounding *wtf* genes. (A) An inversion whose breakpoints fall into the gene bodies of *wts22* and *wts23* in JB900. This inversion was likely caused by recombination between these two *wtf* genes. (B) An inversion altering the gene order surrounding *wts13* in JB872. This inversion was likely caused by recombination between solo LTRs. (C) An inversion altering the gene order surrounding *wts9* in JB943 and JB872. This inversion was likely caused by recombination between full-length Tf transposons. (D) A translocation altering the gene order surrounding *wts7* in JB1180. This translocation was likely caused by recombination between full-length Tf transposons. (E) Duplication of *wts23* in JB1197. This duplication was likely caused by recombination between solo LTRs.

**Supplementary Figure 3.**
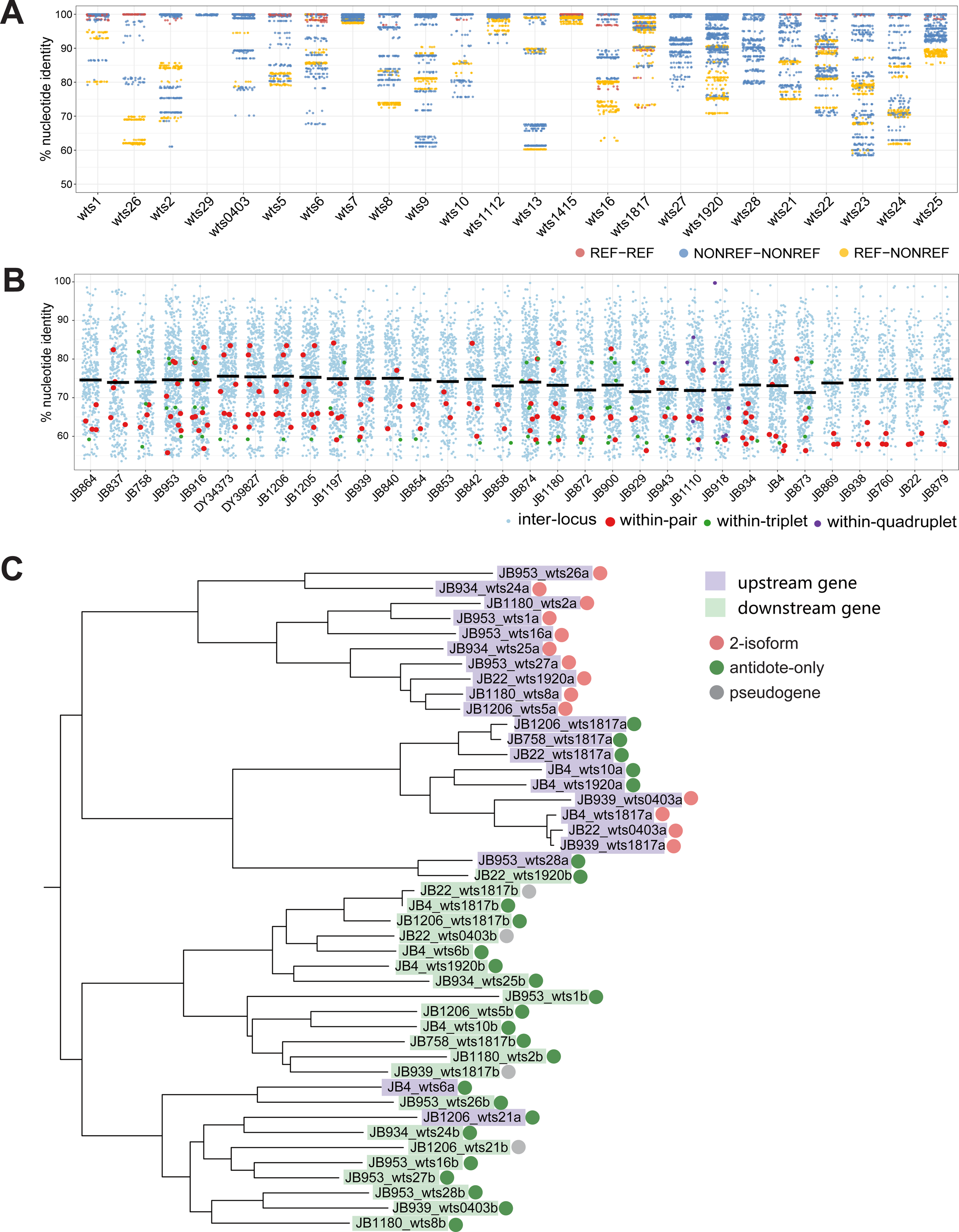
Sequence diversity of *wtf* genes. (A) Pairwise nucleotide identities of syntenic *wtf* genes in different isolates (inter-isolate comparisons). For genes located in multi-gene loci, comparisons were only made when the syntenic loci contained the same number of *wtf* genes. Three different colors are used to distinguish comparisons between genes sharing the REF-lineage ancestry, between genes sharing the NONREF-lineage ancestry, and between genes having different lineage ancestry. Genes in loci with uncertain ancestry are not included in this analysis. (B) Pairwise nucleotide identities of typical *wtf* genes in the same isolate. The median identities are represented by black horizontal lines. Different colors are used to distinguish comparisons between genes in different loci (inter-locus comparisons) and comparisons between genes in the same multi-gene locus (within-pair, within-triplet, and within-quadruplet comparisons). (C) Phylogenetic relationship of 44 *wtf* genes from 22 representative gene pairs. These gene pairs were selected to maximize diversity. The gene names are shown with backgrounds of different colors to distinguish between upstream genes and downstream genes. Color-filled circles next to the gene names denote the functional type of genes (2-isoform, antidote-only, or pseudogene). The phylogenetic tree was manually rooted at the branch separating the two clusters to which most upstream genes and most downstream genes respectively belong.

**Supplementary Figure 4.**
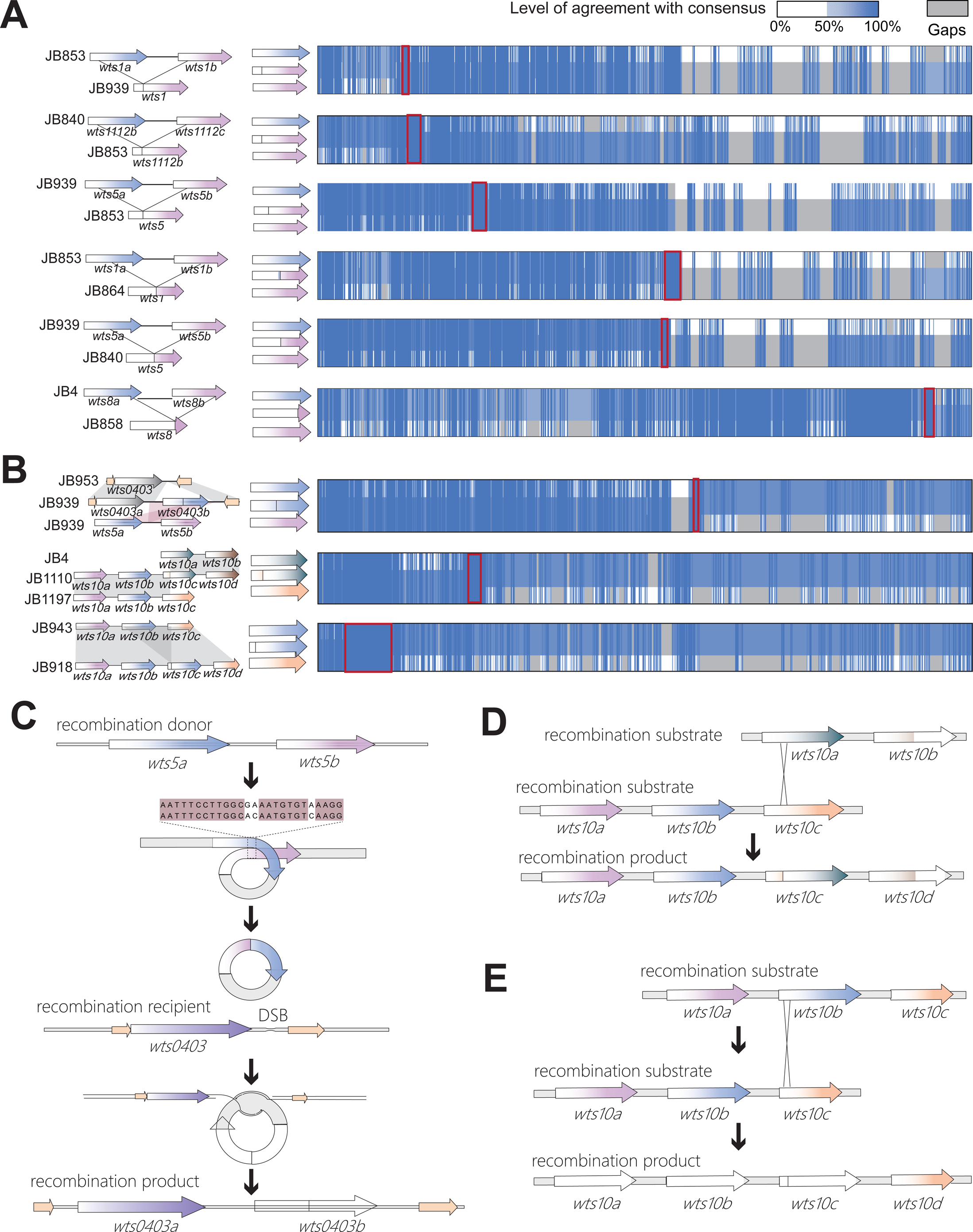
Recent evolutionary events altering the number of *wtf* genes per locus. (A) The 6 events decreasing the number of *wtf* genes per locus. Each event is illustrated with a schematic on the left side, depicting two *wtf* genes in the ancestral allele in a representative isolate (top) and a *wtf* gene resulting from recombination of the two ancestral genes in the derived allele in a representative isolate (bottom). The two representative isolates in each case belong to the same phylogroup. The functional types of the genes are depicted using color-filled circles, as shown in Figure 2. The Jalview-generated alignment overview shown on the right side provides more details on how recombination occurs, with the recombination breakpoint highlighted by a red box. (B) The 3 events increasing the number of *wtf* genes per locus. The schematics on the left side are as follows: for the first event, the ancestral allele (top), the recombination product in the derived allele (middle), and the recombination donor (bottom) are shown; for the second event, two ancestral alleles are shown at the top and bottom, respectively, and the derived allele resulting from recombination of the two ancestral alleles is shown in the middle; for the third event, the ancestral allele is shown at the top, and the derived allele resulting from recombination of two copies of the ancestral alleles is shown at the bottom. The functional types of the genes are depicted using color-filled circles as in Figure 2. The Jalview-generated alignment overview shown on the right side provides more details on how recombination occurs, with the recombination breakpoint highlighted by a red box. (C-E) Schematics illustrating the detailed processes of the events shown in (B).

**Supplementary Figure 5.**
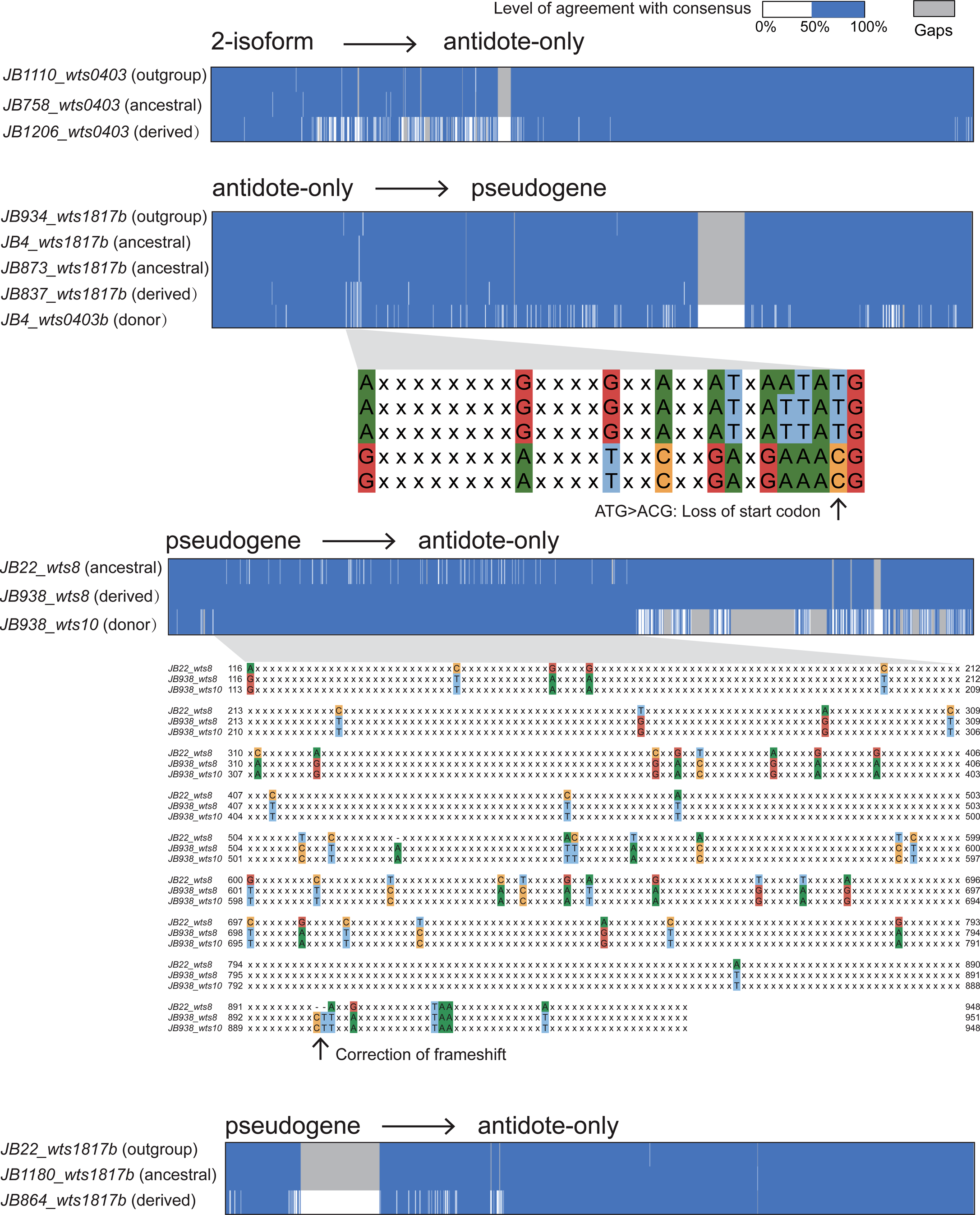
Non-allelic gene conversion events changing the functional type of *wtf* genes. Jalview-generated alignment overviews illustrate where gene conversion occurred. In the case of two events, the nucleotide sequences in the regions that underwent gene conversion are shown in zoomed-out views of the alignments. In these zoomed-out views, identical nucleotides are represented as “x”.

**Supplementary Figure 6.**
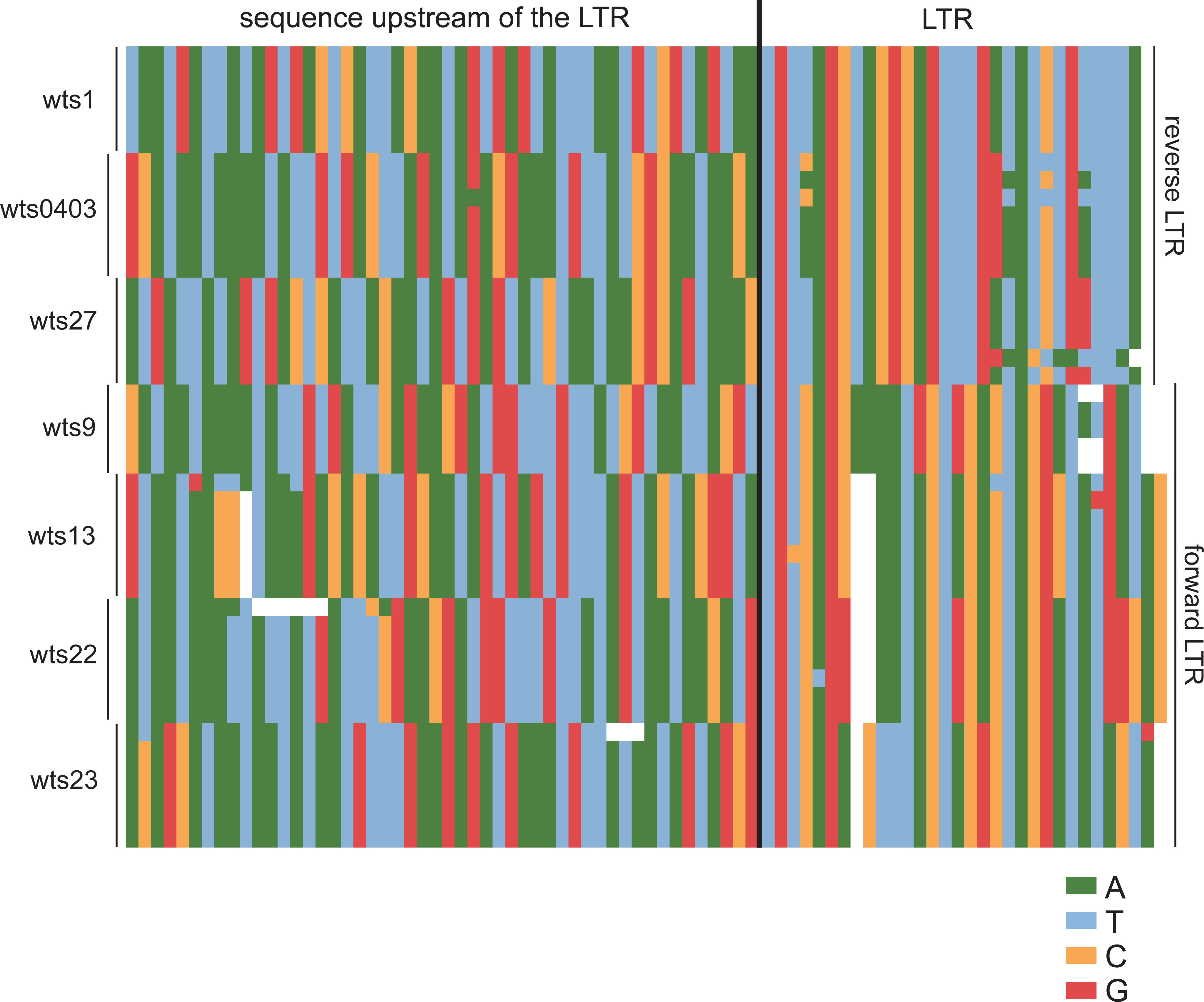
Lack of inter-locus similarities among sequences on the *wtf*-gene-proximal side of the LTRs located downstream of a *wts* locus. In the nucleotide sequence alignment, each sequence includes a 50-bp sequence upstream of the LTR and a 30-bp *wtf*-gene-proximal portion of the LTR. The sequences are from 7 representative isolates and are ordered from top to bottom as follows: JB22, JB1180, JB4, JB758, JB864, JB953, and JB872. For the *wts27* locus, there are only 6 sequences as JB22 does not harbor *wtf* genes at this locus. For the *wts1* locus, there are only 6 sequences due to the exclusion of the sequence JB758, in which an LTR is inserted within the 50-bp sequence. For the *wts9* locus, there are only 5 sequences due to the exclusion of the sequences in JB22 and JB872, because an LTR is inserted within the 50-bp sequence in JB22 and a full-length Tf transposon is inserted within the 50-bp sequence in JB872.

**Supplementary Figure 7.**
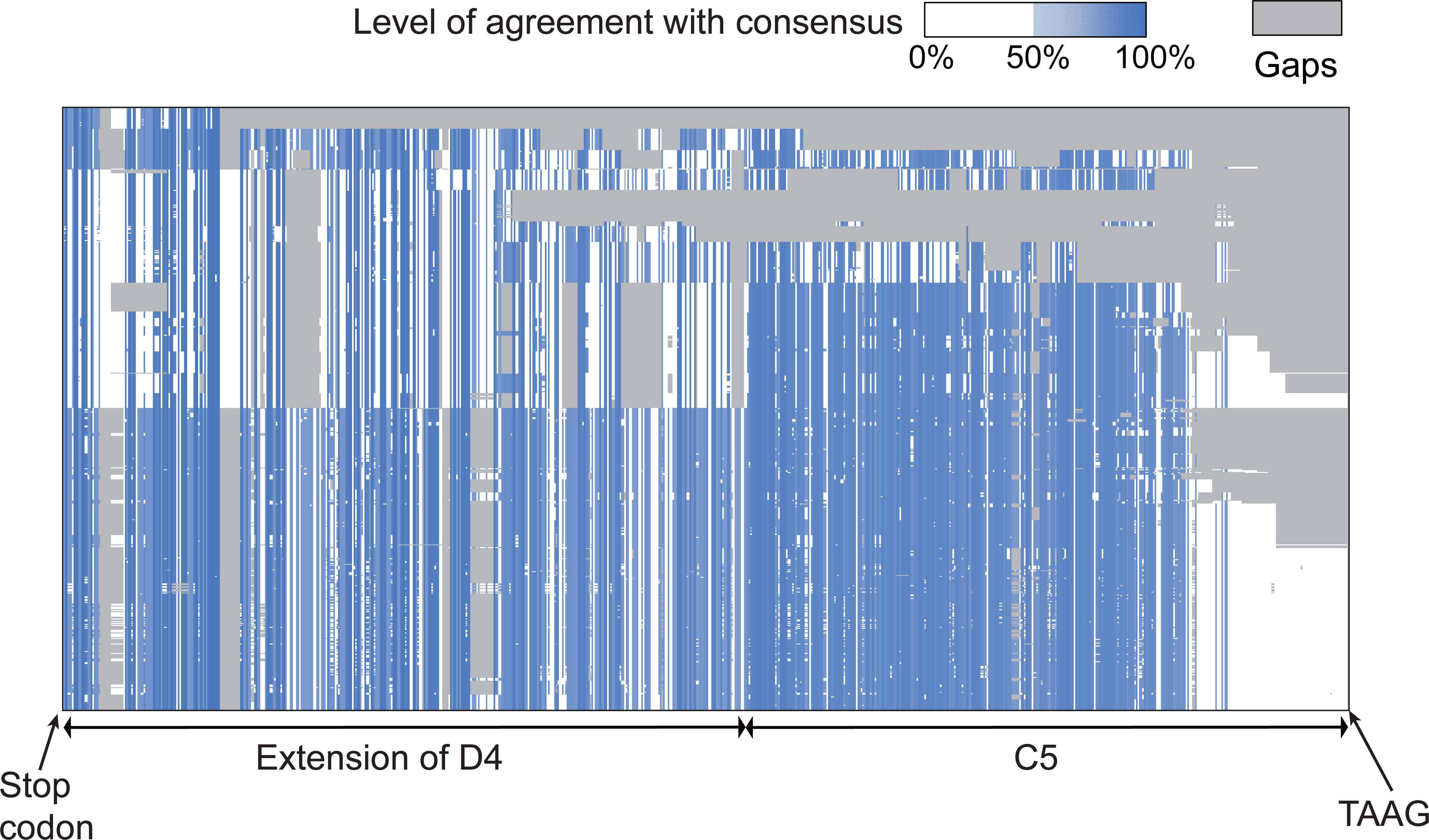
Alignment of the conserved_down sequences. The alignment is shown as a Jalview-generated overview. Gaps in the alignment are shown in dark grey. Nucleotides are colored according to the percentage agreement with the consensus.

**Supplementary Figure 8.**
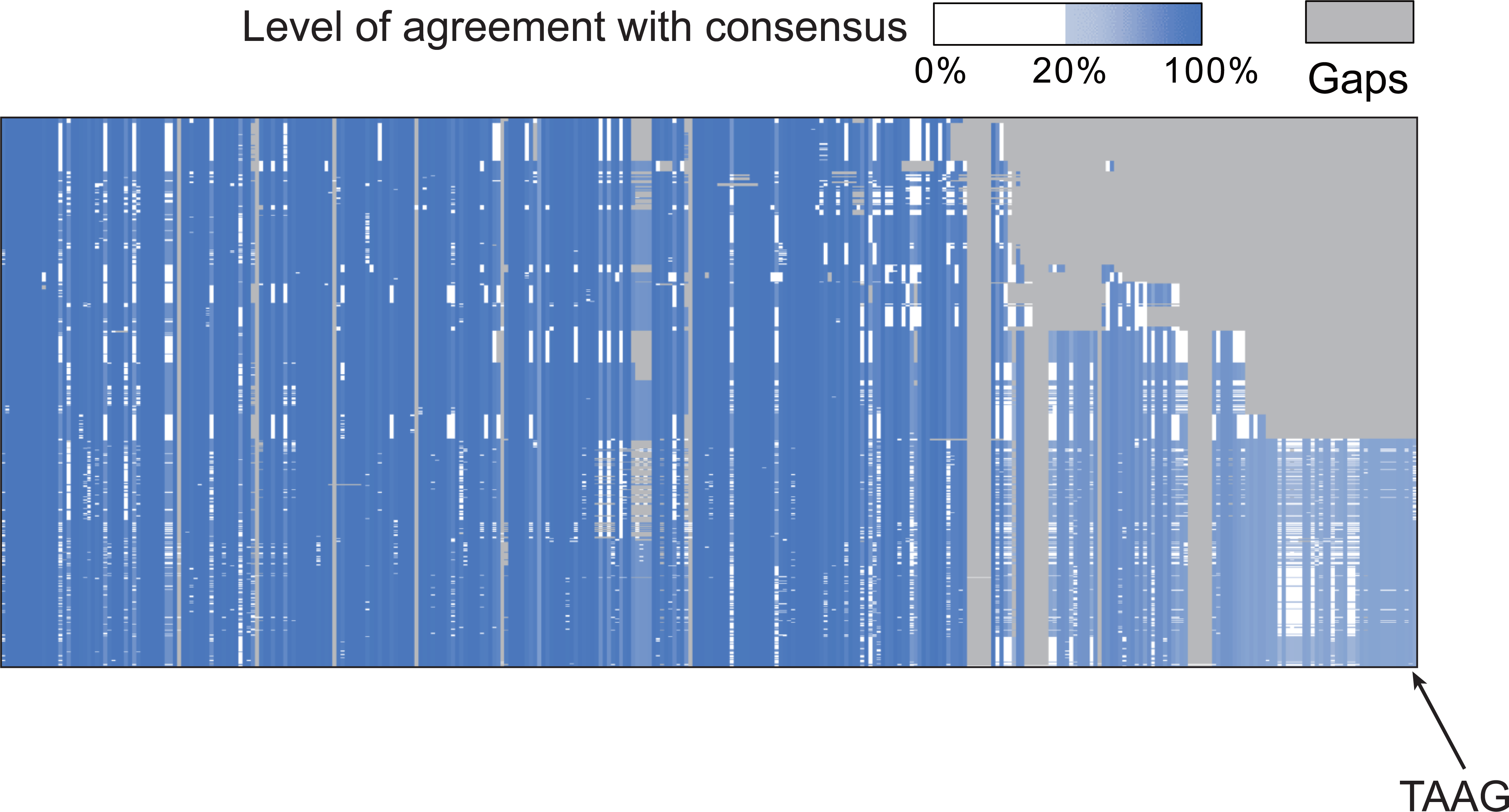
Alignment of the C5 region of conserved_down sequences. The alignment is shown as a Jalview-generated overview. Gaps in the alignment are shown in dark grey. Nucleotides are colored according to the percentage agreement with the consensus.

**Supplementary Figure 9.**
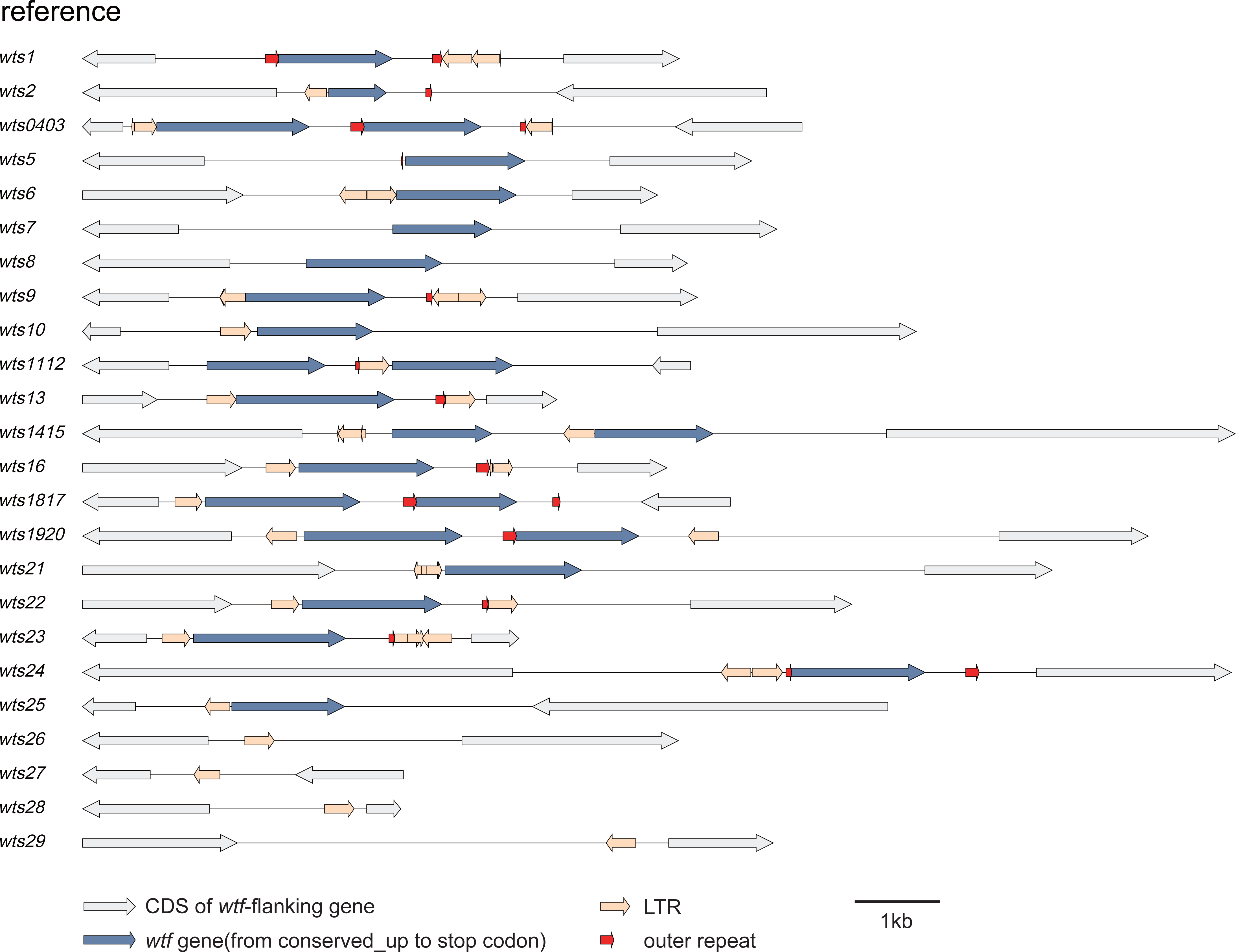
Diagram depicting the outer repeats and *wtf*-flanking LTRs in the reference genome, which is representative of the genomes of REF lineage isolates.

**Supplementary Figure 10.**
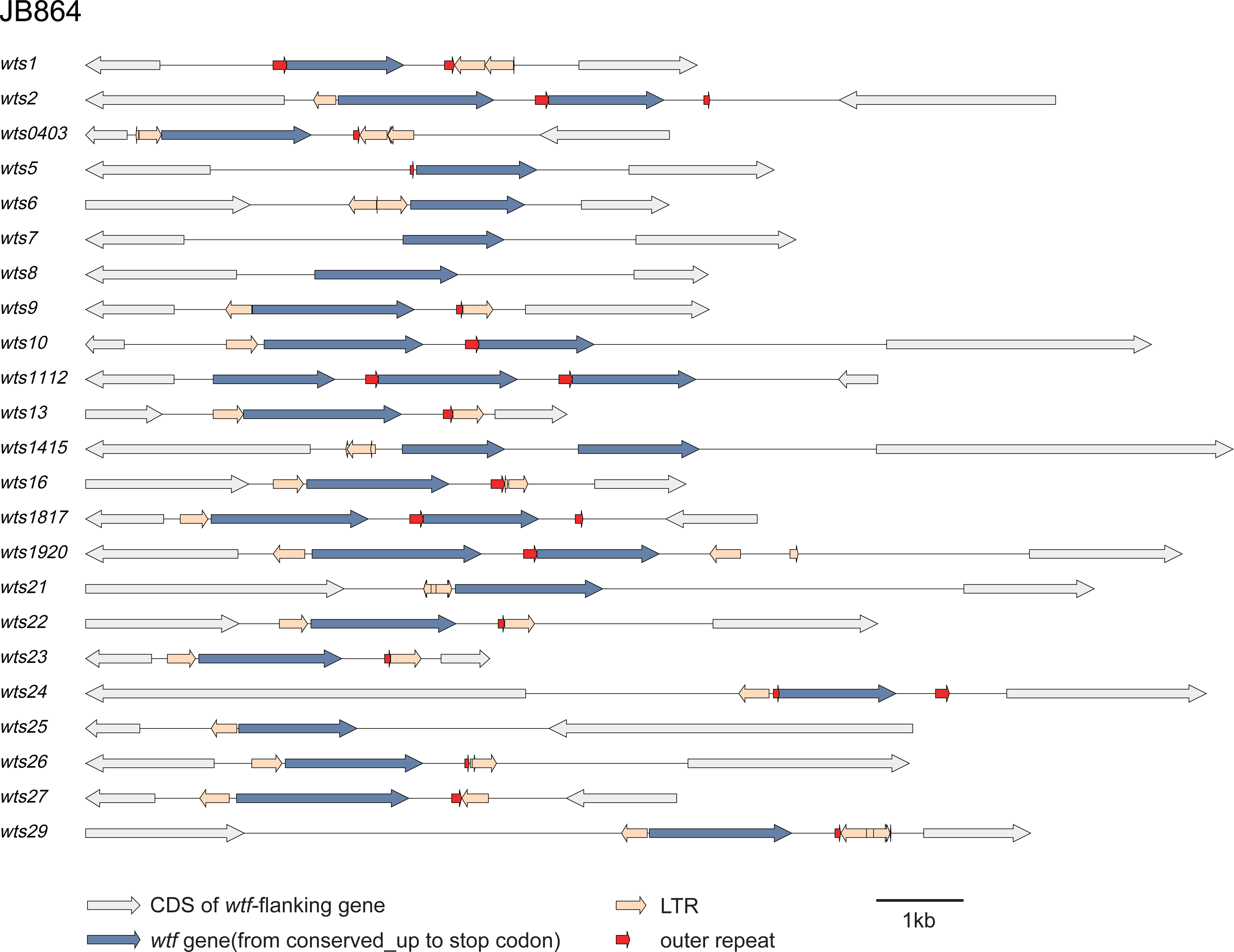
Diagram depicting the outer repeats and *wtf*-flanking LTRs in the genome of JB864, which is representative of the genomes of NONREF lineage isolates. The *wts28* locus is not depicted because the exact sequence of the *wtf* gene at that locus is uncertain.

## Notes

### Competing Interest Statement

The authors have declared no competing interest.

